# Enhancing gene co-expression network inference for the malaria parasite *Plasmodium falciparum*

**DOI:** 10.1101/2023.05.31.543171

**Authors:** Qi Li, Katrina A Button-Simons, Mackenzie AC Sievert, Elias Chahoud, Gabriel F Foster, Kaitlynn Meis, Michael T Ferdig, Tijana Milenković

## Abstract

**Background:** Malaria results in more than 550,000 deaths each year due to drug resistance in the most lethal *Plasmodium* (*P*.) species *P. falciparum*. A full *P. falciparum* genome was published in 2002, yet 44.6% of its genes have unknown functions. Improving functional annotation of genes is important for identifying drug targets and understanding the evolution of drug resistance.

**Results:** Genes function by interacting with one another. So, analyzing gene co-expression networks can enhance functional annotations and prioritize genes for wet lab validation. Earlier efforts to build gene co-expression networks in *P. falciparum* have been limited to a single network inference method or gaining biological understanding for only a single gene and its interacting partners. Here, we explore multiple inference methods and aim to systematically predict functional annotations for all *P. falciparum* genes. We evaluate each inferred network based on how well it predicts existing gene-Gene Ontology (GO) term annotations using network clustering and leave-one-out cross-validation. We assess overlaps of the different networks’ edges (gene co-expression relationships) as well as predicted functional knowledge. The networks’ edges are overall complementary: 47%-85% of all edges are unique to each network. In terms of accuracy of predicting gene functional annotations, all networks yield relatively high precision (as high as 87% for the network inferred using mutual information), but the highest recall reached is below 15%. All networks having low recall means that none of them capture a large amount of all existing gene-GO term annotations. In fact, their annotation predictions are highly complementary, with the largest pairwise overlap of only 27%. We provide ranked lists of inferred gene-gene interactions and predicted gene-GO term annotations for future use and wet lab validation by the malaria community.

**Conclusions:** The different networks seem to capture different aspects of the *P. falciparum* biology in terms of both inferred interactions and predicted gene functional annotations. Thus, relying on a single network inference method should be avoided when possible.

**Supplementary data:** Attached.

**Availability and implementation:** All data and code are available at https://nd.edu/~cone/pfalGCEN/.

## Introduction

### Motivation and related work

Malaria is a deadly disease caused by protozoan parasites of the genus *Plasmodium* (*P*.) that are transmitted by the bite of female mosquitoes [50, 65, 67]. The most deadly malaria species *P. falciparum* causes more than 0.5 million deaths annually, mostly among children under five years old [16, 51, 52, 77]. Sub-Saharan Africa accounts for 79.4% of malaria cases and 87.6% of deaths [51, 74]. *P. falciparum* has evolved resistance to all antimalarial drugs, making treatment difficult in areas where multidrug resistant parasites are common [13, 26, 30, 36, 42]. The *P. falciparum* research community has developed tools to quickly identify mutations that can be used as markers for drug resistance and genes that are under selection. For some drugs a causal gene has been identified that is the main driver of drug resistance [25]. However, understanding how mutations in that gene confer resistance is a more difficult problem to solve. Unfortunately, 44.6% of genes in the *P. falciparum* genome have unknown function. This lack of knowledge of gene function represents a key challenge to a better understanding of how mutations confer drug resistance. In particular, Gene Ontology (GO) annotations that are often used to describe the biology (e.g., biological processes, molecular-level activities, or cellular structures and localization) of genes are deficient for *P. falciparum* [49]. Deriving novel gene-GO term associations would be a valuable contribution to *P. falciparum* gene annotations.

Because biological functions that lead to key traits like drug resistance are controlled by many interacting genes, studying them as complex networks of gene-gene (or protein-protein) interactions presents promising analytical approaches to uncovering important *P. falciparum* biology [23]. Despite pioneering efforts to obtain physical protein-protein interaction (PPI) data for *P. falciparum* [18, 34], high-quality data of this type are lacking [55]. This increases the urgency to understand other interaction/network types. Fortunately, a wealth of gene expression data is available for *P. falciparum* [21], from which gene co-expression networks can be constructed. Gene co-expression networks can be powerfully applied because genes that are functionally related (i.e., that are annotated by the same GO terms) are likely to be co-expressed [73]. Consequently, analyses of a gene co-expression network, where nodes are genes and edges are co-expression relationships between genes over different conditions (e.g., time points or drug treatments), is a valuable tool for identifying novel (i.e., currently unknown) gene-GO term annotations.

Earlier efforts to build gene co-expression networks in *P. falciparum* have been limited to a single network inference method such as mutual information (MI) [60] or absolute value of the Pearson Correlation Coefficient (absPCC) [6, 64], leaving other potentially powerful gene co-expression network inference approaches unexplored. Prominent examples include a tree-based measure called Random Forest (RF) [24] and Adaptive Lasso (AdaL) [32]. In fact, it was shown in different species (baker’s yeast, brewer’s yeast, *E. coli* and *Staphylococcus aureus*) that networks resulting from different gene co-expression network inference methods may be able to give insights into different biological questions [56] and capture different types of regulatory interactions [43]. In addition to investigating *P. falciparum* biology utilizing a single network inference method to build gene co-expression networks, previous efforts have asked limited biological questions about the interacting partners or functions of particular genes [1, 21, 60, 66, 79]. However, gene co-expression networks developed using multiple inference methods have not yet been developed for *P. falciparum*. It is unknown whether different network inference methods capture different aspects of *P. falciparum* biology and whether they could inform the important task of systematically predicting GO term annotations for all *P*.*falciparum* genes.

Here, we fill these gaps by constructing multiple co-expression networks using four prominent network inference methods (MI, absPCC, RF, and AdaL) and evaluate them for systematic, comprehensive, and accurate predictions of gene-GO term associations. Furthermore, we assess the extent to which the networks are complementary or redundant in terms of their edges (i.e., gene co-expression relationships) as well as predicted functional knowledge. Finally, we apply our inferred networks to the study of endocytosis, an essential biological process involved in *P. falciparum* drug resistance.

### Our study and contributions

We analyze gene expression data consisting of 247 samples corresponding to 247 combinations of drug treatments and time points [21] (Fig. 1). We apply four network inference methods to this expression data to infer their respective co-expression networks. Each network inference method mathematically assigns a weight representing the strength of a co-expression relationship between a pair of genes. The highest-weighted gene (i.e., node) pairs are kept as edges in the given network. To choose a weight threshold for distinguishing between edges and non-edges in a network, we adopt a prominent network inference framework called ARACNe [44].

**Figure 1.**
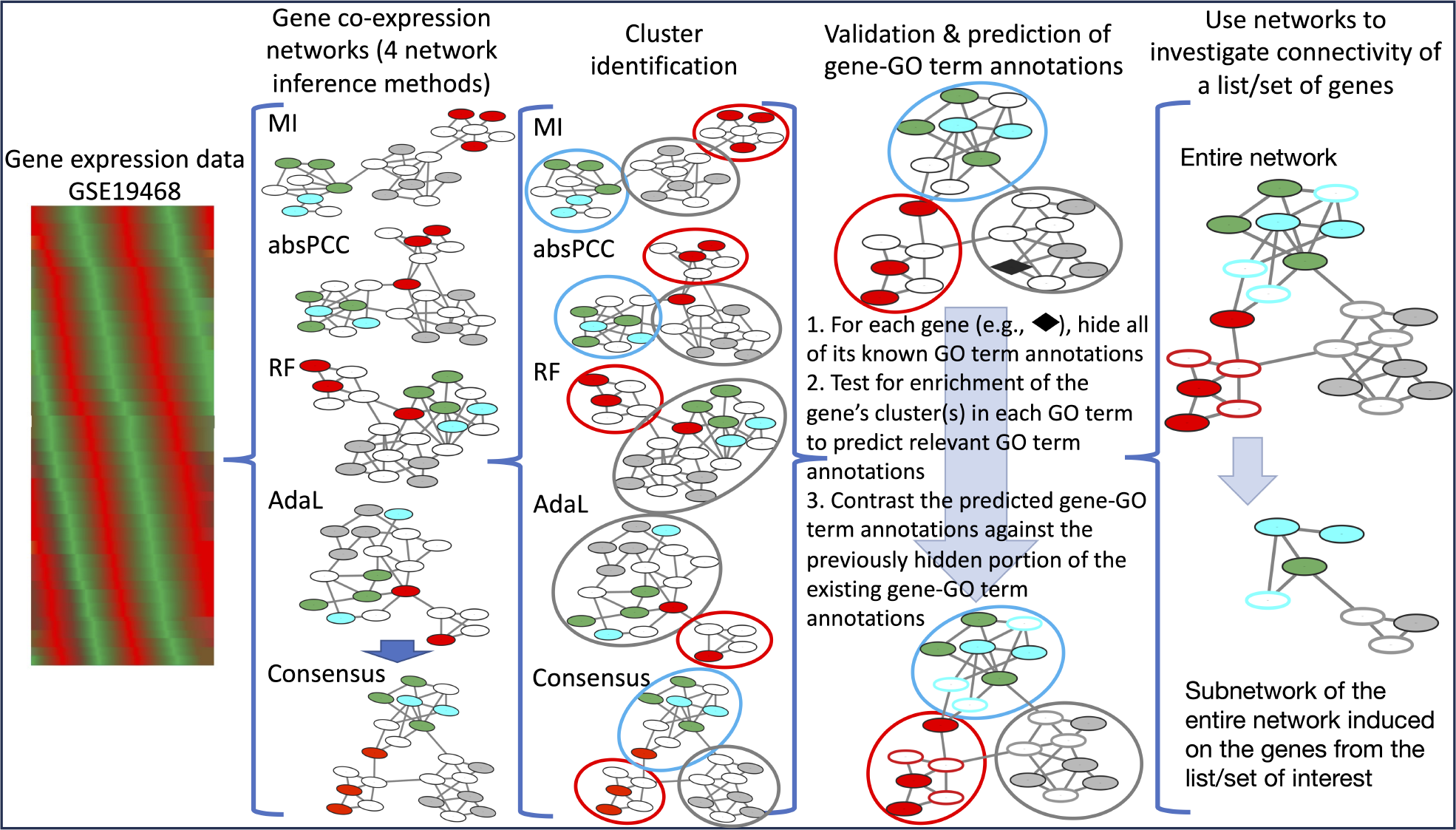
Summary of our generalizable framework for gene co-expression network construction, validation, and biological application. We use four network inference methods (MI, absPCC, RF, and AdaL) to construct gene co-expression networks of *P. falciparum*; note that in the illustration, nodes with the same color indicate that they are annotated by the same GO term. We also generate a Consensus network by combining the four individual networks. We use two clustering methods to identify gene clusters (functional modules) in each network. Then, we use cross-validation to examine how well each network’s clusters correspond to existing GO terms. In other words, GO term annotations are predicted for all genes whose clusters are statistically significantly enriched in at least one GO term, and the predicted gene-GO term annotations are then contrasted against a previously hidden portion of the existing ones. The Consensus network is used to investigate connectivity of a gene list/set; in our biological application, we investigate lists of genes that interact with three proteins involved with the biological process of endocytosis.

We find that the co-expression networks resulting from the different inference methods are overall highly complementary (i.e., non-redundant). Namely, the networks constructed using MI, absPCC, and AdaL, share only 15%-53% of the edges; that is, 47%-85% of the edges are unique to each of the networks. Only the network resulting from RF is mostly redundant to the networks resulting from the other three inference methods. This indicates that the different network inference methods largely capture unique features of gene co-expression relationships. Given so many unique edges in almost each of the networks, we consider an additional co-expression network that integrates (takes union of) the edges from all of the individual networks; we refer to this network as Consensus.

Consequently, we use gene-GO term annotations (or associations) to assess each co-expression network by leveraging data from GeneDB ^1^ and PlasmoDB ^2^ databases. We hide a portion of the existing (i.e., ground truth) gene-GO term annotations (or associations), use the remaining non-hidden associations along with the given network’s structure (or topology) to predict additional gene-GO term annotations, and evaluate how well the predicted annotations match the hidden ones, all using cross-validation. The better this match, the more functionally meaningful the given network’s topology, i.e., the better the corresponding network inference method.

Specifically, we use a common paradigm for predicting function annotations called network clustering. A cluster is a group of genes in a network that are densely connected to each other or have similar topological patterns [**?**]. A cluster is deemed functionally meaningful if a statistically significantly high number of its genes are annotated by the same GO term. By extension, one can then predict the other genes in this cluster to also be annotated by this GO term. We use two clustering methods (i.e., cluster affiliation model for big networks – BigCLAM, referred to as BC [78], and Markov Clustering – referred to as MCL [14]) as well as multiple parameter values for each clustering method. We use these clustering methods because they are highly prominent [15, 40, 41, 47, 53, 72], and also, MCL has been shown to perform consistently well in different contexts [11, 40, 47, 71, 72].

We cluster each network and predict gene-GO term annotations from each of the given network’s clusters. We evaluate a network’s prediction accuracy over all of its clusters via measures of precision and recall, by comparing the predicted annotations with the hidden portion of the existing gene-GO term annotations. Precision is the fraction of the predicted annotations that are correct (i.e., that currently exist); recall is the fraction of all existing annotations that are predicted. Generally there is a trade-off between precision and recall: higher precision typically means lower recall, and vice versa. In biomedical applications, precision is typically favored over recall [31, 35, 38, 39, 48] because confirmatory wet lab experiments are time consuming and expensive and it is often preferred to follow-up on a few higher-quality predictions than many lower-quality ones.

In our own analysis, while some of the networks generate precision scores as high as 87%, the highest recall reached is below 15%. The MI method generates the highest precision and the others also have reasonably high precision scores. All networks have relatively low recall indicating that none capture a large amount of all existing gene-GO term annotations. In fact, we find the different networks’ predictions to be highly complementary to each other (i.e., the maximum prediction overlap over all network pairs is only 27%). Hence, the different networks seem to capture different aspects of the *P. falciparum* biology, as was the case in baker’s yeast [56].

To supplement the limited functional annotation data in the *P. falciparum* genome, we assign a confidence score to each (existing, i.e., currently known, and novel, i.e., currently unknown) predicted gene-GO term association based on how many networks support the given prediction. Similarly, we assign confidence scores to the inferred gene-gene interactions, i.e., co-expression relationships. We provide the confidence score-ranked lists of functional predictions and interactions for future community use that could be especially useful given the paucity of comprehensive functional annotations as well as high confidence PPI data for *P. falciparum*.

In addition to validating inferred networks via gene functional prediction, we investigate the connectivity of genes hypothesized to function together in a biological process. That is, we apply our network approach to recently generated lists of endocytosis-related genes in *P. falciparum* [7]. Endocytosis is an essential biological process for trafficking extracellular material to specific organelles in the cell. Extracellular material is first brought into the cell by early endosomes that mature into late endosomes as they are directed to lysosomes or through the trafficking pathways from the Golgi apparatus. The *P. falciparum* proteins Kelch13 (K13) and EPS15 have been localized to the periphery of the cell and suggested to participate in the machinery to generate early endosomes. The *P. falciparum* protein clathrin functions in an atypical role, primarily through trafficking pathways of the Golgi apparatus in other apicomplexan organisms with similar functions suggested in *Plasmodium* [20, 54]. K13 is a molecular determinant of artemisinin resistance. EPS15 is a close interacting partner of K13 and typically functions in canonical endocytosis pathways [68], and clathrin is also the main structural protein in clathrin-dependent endocytosis. Recently, Birnbaum *et al*. [7] experimentally determined lists of genes that interact via PPIs with K13, EPS15, and clathrin. Their K13 and ESP15 gene lists were devoid of clathrin, suggesting that K13 functions in a clathrin-independent endocytosis pathway. To explore these and other proposed interactions, we use our Consensus co-expression network data for an independent systems-level view as a way to generate new hypotheses about this cellular pathway and the mechanism of resistance for artemisinin. For each of the K13, EPS15, and clathrin gene lists, we find that the genes in a given list are statistically significantly more densely interconnected with each other in the Consensus network than expected by chance, supporting and extending the claim that these genes are involved in the same biological process, endocytosis. We also validate some additional hypotheses of Birnbaum *et al*. [7], which further validates our inferred network data, while also providing additional novel insights into the roles of K13, EPS15, and clathrin in *P. falciparum* endocytosis pathways.

## Results and discussion

### Description of and overlap between inferred networks

We consider four network inference methods to infer respective co-expression networks from gene expression data GSE19468 containing 4,374 genes (Section *“Inference of gene co-expression networks”*, also Additional File 6). Each network is named based on its inference method, i.e., MI, absPCC, RF, and AdaL. Among these, as presented in Table 1, all four networks have a similar number of nodes, ranging between 4,082 and 4,374. However, the number of edges and thus network density (the percentage of edges that exist out of all possible edges) varies drastically across the four networks. The one with the lowest network density is AdaL (0.09%), followed by MI (0.77%), then RF (9.08%), and finally absPCC (10.19%).

**Table 1.** The 11 considered co-expression networks and their size and density statistics.

| Network | # of nodes | # of edges | Density |
| --- | --- | --- | --- |
| MI | 4,373 | 73,474 | 0.77% |
| absPCC-0.3 | 3,792 | 285,405 | 3.97% |
| absPCC-0.4 | 3,993 | 380,539 | 4.77% |
| absPCC-0.5 | 4,126 | 475,674 | 5.59% |
| absPCC | 4,322 | 951,347 | 10.19% |
| RF-0.03 | 4,188 | 25,000 | 0.29% |
| RF-0.05 | 4,323 | 43,406 | 0.46% |
| RF-0.1 | 4,372 | 86,811 | 0.91% |
| RF | 4,374 | 868,105 | 9.08% |
| AdaL | 4,082 | 7,708 | 0.09% |
| Consensus | 4,374 | 333,162 | 3.48% |

A smaller density indicates a more conservative network inference method, i.e., a method that judges fewer co-expressions as strong enough. Two networks, absPCC and RF, exhibit extremely higher densities than typically sparse real-world networks, including PPI and other biological networks [4]. To handle these two methods being unusually non-conservative, i.e., to highlight the most important gene co-expression relationships within absPCC and RF, as is typically done [**?**], we examine whether we can remove some proportion of the edges without disconnecting many of the nodes from the given network, as such edges could be viewed as redundant. We do this systematically. Namely, for each of the two networks, we keep 1%-100% (in increments of 2% between 1%-50% and increments of 5% between 50%-100%) most important (highest weighted) edges among all of the edges in a given network. Then, for each of these thresholds, we calculate the percentage of nodes from the gene expression data that are in the largest connected component of the resulting thresholded network.

We aim to preserve at least 85% of the genes from the gene expression data in the largest connected component. This corresponds to keeping at least 30% most important edges in the absPCC network. So, we consider the threshold value of 30%, along with two additional, arbitrarily chosen higher values, namely 40% and 50%, for further systematic evaluation. We refer to these three absPCC-based subnetworks as absPCC-0.3, absPCC-0.4, and absPCC-0.5, respectively. Also, from the systematic thresholding procedure described above, we find an interesting pattern with the RF network. Namely, keeping as few as 3% most important edges already results in more than 95% of the genes from the gene expression data being in the largest connected component. This is why we select this threshold value of 3%, along with two additional, arbitrarily chosen higher values of 5% and 10%. We refer to these three RF-based subnetworks as RF-0.03, RF-0.05, and RF-0.1, respectively. Supplementary Fig. S1 further illustrates the rationale for determining the number of edges kept. Therefore, up to this point, we have constructed 10 networks, one inferred using MI, four inferred using absPCC, four inferred using RF, and one inferred using AdaL (Table 1; note that this table contains an additional network, Consensus, which is discussed below).

These results indicate that the complementary co-expression networks may be largely capturing different aspects of *P. falciparum* biology. This is exactly what we find in our subsequent analyses discussed below, in which we show that the networks are predicting complementary gene-GO term annotations. This observation is consistent with results from previous related studies on other systems [43, 56]. Given such high network complementarity, in hope of more accurately capturing regulatory relationships between genes, as is sometimes done [43], we construct a Consensus network by taking the union of all edges from the complementary co-expression networks. For this, because the different network inference methods have different conservativeness levels judging whether an edge is strong enough, i.e., because they possibly weigh the same edge differently, we align these levels between the four networks using min-max normalization. That is, for each network, we normalize the edge weights in a given network to the (0, 1] range. Then, for each edge, we sum its normalized edge weights over the four networks. As such, the resulting Consensus network has edge weights in the (0, 4] range; the higher the weight of an edge, the more important the edge is or the more networks are supporting this edge (or both) (Section *“Methods”*). Therefore, in total, we consider 11 co-expression networks in this study (Table 1).

### Selecting the best clustering parameter values for the inferred networks

We use two clustering methods (i.e., BigCLAM (BC) and MCL) to generate clusters for predicting gene-GO term associations from each of the 11 co-expression networks. For unbiased and comprehensive evaluation of each network and fair comparison of the different networks, we test multiple parameter values for each clustering method in each network. We use three criteria to select (up to) three clustering parameter values: i) the parameter value that yields the highest precision in cross-validation; ii) the parameter value for which the union of all clusters that are significantly enriched in one or more GO terms contains the most of unique genes, i.e., has the largest gene coverage; and iii) the parameter value for which the union of all clusters that are significantly enriched in one or more GO terms contains the most of unique GO terms, i.e., has the largest GO term coverage.

Intuitively, for each combination of a network, a clustering method, and a clustering parameter value, we obtain a set of clusters. We use leave-one-out cross-validation to predict gene-GO term association from a given set of clusters. That is, given a gene *g*, we hide its existing GO term annotations. Then we examine whether each of the clusters that gene *g* belongs to is statistically significantly enriched (with adjusted *p*-value ¡ 0.05) in an existing GO term *j* using the hypergeometric test; we do this for each GO term under consideration. If so, for such a cluster and GO term *j*, we predict gene *g* being annotated by GO term *j*. We iterate the above process over all genes in the gene expression data and calculate precision and recall, along with the gene coverage and GO term coverage (as defined above) of the significantly enriched clusters. The former serves our criterion i) above, and the latter two serve our criteria ii) and iii) above, respectively. We carry out this entire procedure for each cluster set, i.e., each combination of network, clustering method, and parameter value. Heuristically, these three criteria maximize the accuracy as well as coverage of the predicted gene-GO term annotations for each combination. Note that the three criteria can share the same clustering parameter value (Section *“Methods”*). We list the selected parameter values and their resulting number of clusters in Table 2.

**Table 2.** The best (i.e., selected) clustering parameter value for each of the three criteria, for each of BC and MCL clustering methods, and for each co-expression network. The parameter of BC (whose values range from 25 to 700) corresponds to the expected number of resulting clusters. The parameter of MCL (that starts with an “I” and whose values range from 1.2 to 5) corresponds to the concept of inflation in MCL. Intuitively, the higher the inflation value, the more clusters MCL is expected to return, and hence, the smaller the expected average cluster size is. We list the resulting number of clusters behind each MCL cluster parameter value.

| Network | BC parameters |  |  | MCL parameters |  |  |
| --- | --- | --- | --- | --- | --- | --- |
|  | Criterion i) | Criterion ii) | Criterion iii) | Criterion i) | Criterion ii) | Criterion iii) |
| MI | 425 | 25 | 450 | I2.4 | I1.6 | I1.7 |
| absPCC-0.3 | 350 | 100 | 450 | I1.5 | I2.9 | I2 |
| absPCC-0.4 | 650 | 125 | 500 | I1.6 | I2.3 | I2 |
| absPCC-0.5 | 650 | 275 | 475 | I1.32 | I2.6 | I2.4 |
| absPCC | 650 | 225 | 225 | I3.6 | I2.5 | I3 |
| RF-0.03 | 700 | 25 | 200 | I2.8 | I1.2 | I2 |
| RF-0.05 | 600 | 50 | 350 | I3.2 | I1.24 | I1.9 |
| RF-0.1 | 700 | 50 | 450 | I5 | I1.28 | I2.1 |
| RF | 475 | 50 | 75 | I5 | I1.7 | I2.5 |
| AdaL | 550 | 25 | 75 | I1.24 | I1.2 | I1.7 |
| Consensus | 275 | 275 | 200 | I5 | I1.36 | I1.7 |

According to the selected parameter values, we find that the three criteria result in quite different parameter values, which in turn result in different numbers of clusters and different cluster sizes. This stresses the need of testing multiple parameter values for a given clustering method.

### Validating the inferred networks in the task of predicting gene-GO term associations

For each of the 11 networks, given two clustering methods and three selected parameter values per clustering method, there are six combinations of clustering method and parameter value (Table 2). To further simplify presentation of results, for a given network, we discard from further consideration any of its considered combinations that has both lower precision and lower recall than another one of the considered combinations for the same network. That is, we continue considering only the best combinations, i.e., those combinations that are superior to all other combinations for the same network with respect to at least one of precision and recall. For details, see “*Methods*” and Supplementary Figs. S3–S7. This results in the following combinations for further consideration; for MI network: MCL-I1.6, BC-425, and MCL-I2.4 (Supplementary Fig. S3); for AdaL network: BC-550, MCL-I1.24, and MCL-I1.7 (Supplementary Fig. S6); for Consensus: MCL-I1.7 and MCL-I5 (Supplementary Fig. S7); over all four thresholded absPCC networks: absPCC-BC-650, abcPSS-0.5-BC-650, absPCC-0.3-MCL-I2, absPCC-0.3-MCL-I1.5, and absPCC-0.5-MCL-I3.2 (Supplementary Fig. S4); over all thresholded RF networks: RF-0.1-MCL-I1.28, RF-0.05-BC-600, RF-0.03-BC-700, RF-0.03-MCL-I2, and RF-0.03-MCL-I1.28 (Supplementary Fig. S5).

Given these best combinations of network, clustering method, and clustering parameter value, we present their precision and recall in Fig. 3. We observe that combinations that yield higher precision also yield lower recall, and vice versa. This is expected as there is a trade-off between precision and recall. Precision is typically favored over recall in biomedicine [38, 39]. The recall values of all combinations (and thus of all networks) are below 15%, but some of the networks yield high precision (Fig. 3). In particular, the highest precision value over all combinations for MI is 86.7%, i.e., MI yields precision of 86.7%. It is followed by absPCC-0.5 with precision of 77.8%, RF-0.03 with precision of 74.3%, and AdaL with precision of 56.2%. In other words, with respect to precision, the four top-performing combinations of network, clustering method, and parameter value span all four consideed network inference methods. This means that all inference methods successfully capture meaningful biological signal, i.e., existing gene-GO term annotations. On the other hand, low recall of all combinations (specifically, recall values of 2.2%, 0.4%, 2.5%, and 1.8% corresponding to the above four precision values of 86.7%, 77.8%, 74.3%, and 56.2%, respectively) indicates that none of the network inference methods, i.e., their resulting networks, capture a large fraction of all existing gene-GO term annotations. Therefore, each of the four network inference methods has a value, but none of them is sufficient on its own. This observation aligns with the motivation of inferring the Consensus network in hope to increase both precision and recall of the individual networks.

**Figure 2.**
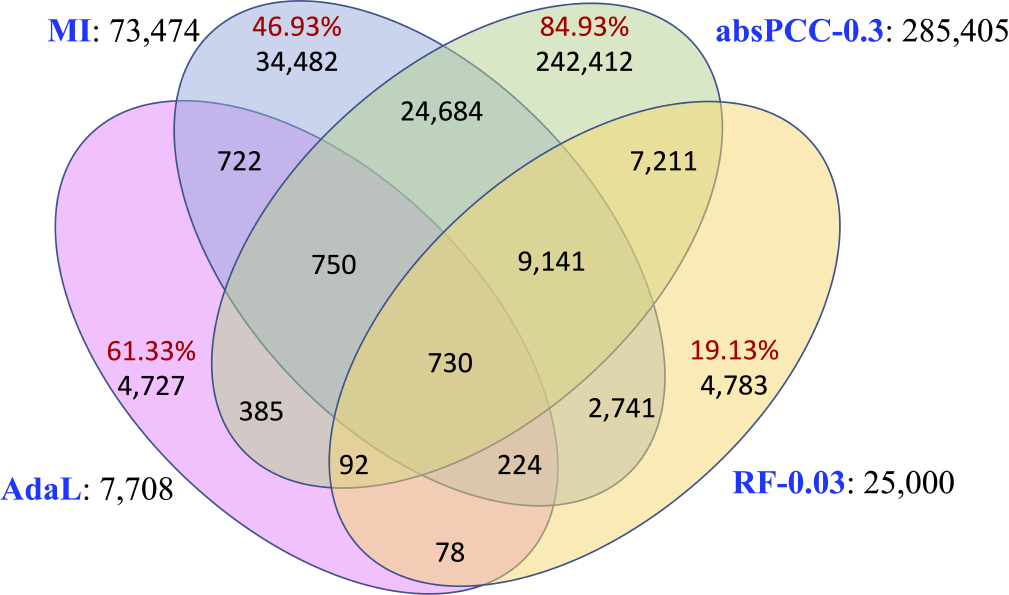
The edge overlaps between the four networks. Each network’s name is colored in blue, and is followed by its corresponding number of edges. Within the Venn diagram, each red number is the percentage of all edges in its corresponding network that are unique to the given network. For example, out of all 7708 edges in AdaL, 4727 (i.e., 61.33% of the) edges are unique to AdaL. More detailed information can be found in Supplementary Fig. S2.

**Figure 3.**
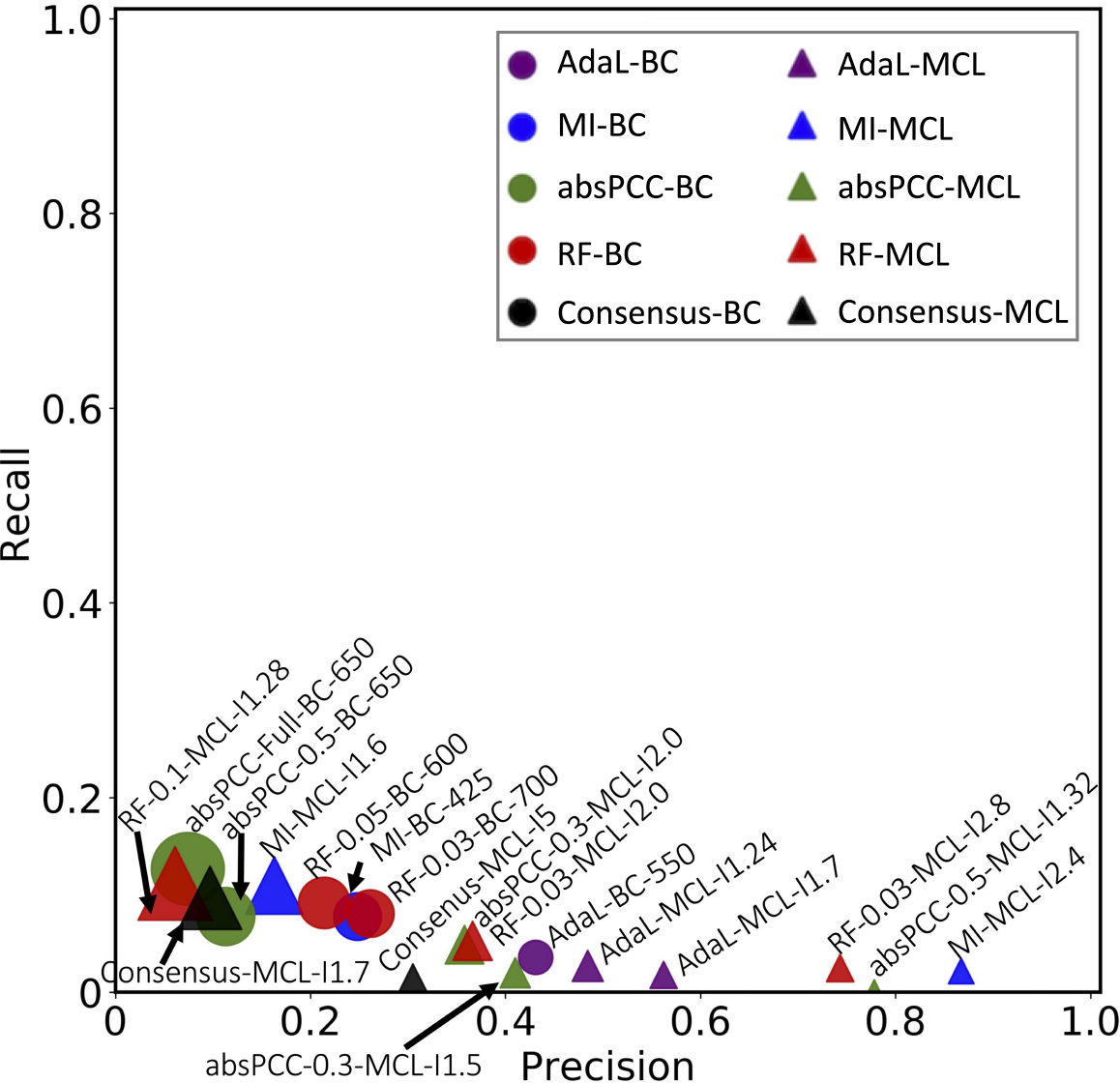
Accuracy of predicting gene-GO term associations in the leave-one-out cross-validation in terms of precision and recall. Each point is a combination of network, clustering method, and parameter value. The sizes of the points correspond to the numbers of predictions produced by a given combination. The color of a point corresponds to a network, and the shape of a point corresponds to a clustering method. For example, all purple points correspond to AdaL, of which circles correspond to BC and triangles correspond to MCL. Note that we use one color (green) for all absPCC networks corresponding to different edge thresholds (e.g., absPCC and absPCC-0.5 share the same color), and we use another color (red) for all RF networks corresponding to different edge thresholds.

However, we find that the Consensus network does not perform the best (Fig. 3). In fact, it performs worse than all four top-performing combinations involving the four individual networks that Concensus is constructed from. This could be because the four individual networks capture complementary edges, and simply combining their edges (even via the edge weighing scheme that we use) into Consensus does not necessarily mean producing more biologically meaningful clusters than in individual networks. This might especially hold if the clusters formed by the edges in one individual network capture different GO terms (i.e., different biological knowledge) than the clusters formed by the edges in another individual network; we explore potential complementarity of the biological knowledge captured by the individual networks later on in this section. If this is the case, integrating the edges from the different networks into the Consensus network and then making functional predictions from this network might weaken the biological signal that can be extracted from the individual networks. But using the individual networks to first make their functional predictions, with each individual network resulting in high precision but low recall, and then integrating the predictions, should result in still high precision, hopefully close to the precision values of the individual networks, but now also higher recall than the recall values of the individual networks. Indeed, we verify that this is what happens: when we integrate functional predictions of the four top-performing (in terms of precision in Fig. 3) combinations of network, clustering method, and parameter value that cover all four network inference methods, the resulting precision is 66% (compared to the individual precision values of 86.7%, 77.8%, 74.3%, and 56.2%) and the resulting recall is 5% (compared to the individual recall values of 2.2%, 0.4%, 2.5%, and 1.8%). Importantly, even though in the task of predicting gene-GO term annotations the Consensus network does not perform better than the individual networks, the Consensus network does successfully capture drug resistance “biology” relevant to endocytosis (Section *“Endocytosis-related biological signatures are captured by the Consensus network”*).

However, we find that the Consensus network does not perform the best (Fig. 3). In fact, it performs worse than all four top-performing combinations involving the four individual networks that Concensus is constructed from. This could be because the four individual networks capture complementary edges, and simply combining their edges (even via the edge weighing scheme that we use) into Consensus does not necessarily mean producing more biologically meaningful clusters than in individual networks. This might especially hold if the clusters formed by the edges in one individual network capture different GO terms (i.e., different biological knowledge) than the clusters formed by the edges in another individual network; we explore potential complementarity of the biological knowledge captured by the individual networks later on in this section. If this is the case, integrating the edges from the different networks into the Consensus network and then making functional predictions from this network might weaken the biological signal that can be extracted from the individual networks. But using the individual networks to first make their functional predictions, with each individual network resulting in high precision but low recall, and then integrating the predictions, should result in still high precision, hopefully close to the precision values of the individual networks, but now also higher recall than the recall values of the individual networks. Indeed, we verify that this is what happens: when we integrate functional predictions of the four top-performing (in terms of precision in Fig. 3) combinations of network, clustering method, and parameter value that cover all four network inference methods, the resulting precision is 66% (compared to the individual precision values of 86.7%, 77.8%, 74.3%, and 56.2%) and the resulting recall is 5% (compared to the individual recall values of 2.2%, 0.4%, 2.5%, and 1.8%). Importantly, even though in the task of predicting gene-GO term annotations the Consensus network does not perform better than the individual networks, the Consensus network does successfully capture drug resistance “biology” relevant to endocytosis (Section *“Endocytosis-related biological signatures are captured by the Consensus network”*).

Next, as mentioned above, we examine whether the four top-performing combinations of network, clustering method, and parameter value that cover all four network inference methods, plus the Consensus network, yield redundant or complementary predicted gene-GO term associations. That is, here, we analyze MI-MCL-I2.4, absPCC-0.5-MCL-I1.32, RF-0.03-MCL-I2.8, AdaL-MCL-I1.7, and Consensus-I5 networks. For each pair of these networks, we measure overlaps of 1) predicted gene-GO term associations, 2) unique genes that participate in the predicted associations, and 3) unique GO terms that participate in the predicted associations. We do all of this with respect to i) predicted existing associations, i.e., true positives (associations that currently exist and are predicted by the networks) (Fig. 4) as well as ii) novel associations (associations predicted by the networks that do not currently exist) (Fig. 5). We quantify the size of an overlap using the Jaccard index, where a lower Jaccard index indicates a lower redundancy, i.e., a higher complementarity.

**Figure 4.**
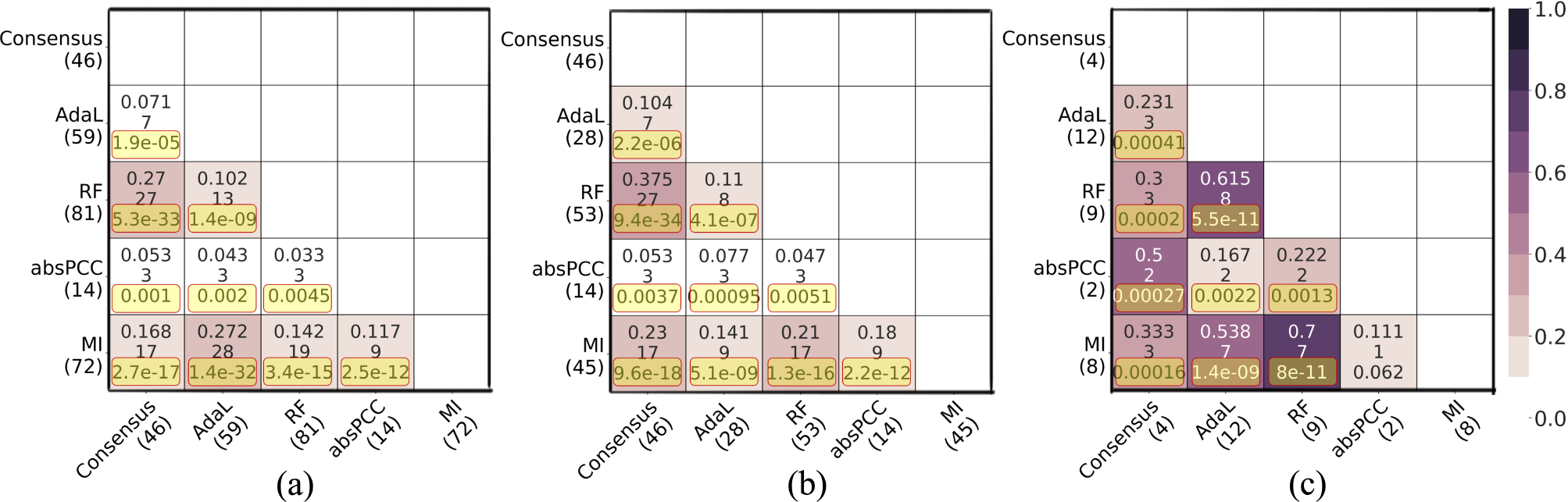
Pairwise overlaps between predictions made by the best combination for each of the five networks with respect to predicted existing associations, i.e., true positives. Panel (a) shows pairwise overlaps of gene-GO term associations. Panel (b) shows pairwise overlaps of unique genes that participate in the predicted gene-GO term associations. Panel (c) shows pairwise overlaps of unique GO terms that participate in the predicted gene-GO term associations. Within each panel, the number in parentheses below each network name is the number of corresponding true positives. Within each cell, there are three numbers; the first number is the Jaccard index, the second number is the raw number of the predictions in the given overlap, and the third number is the adjusted *p*-value resulting from the hypergeometric test. The yellow boxes highlight the adjusted *p*-values that are statistically significant.

**Figure 5.**
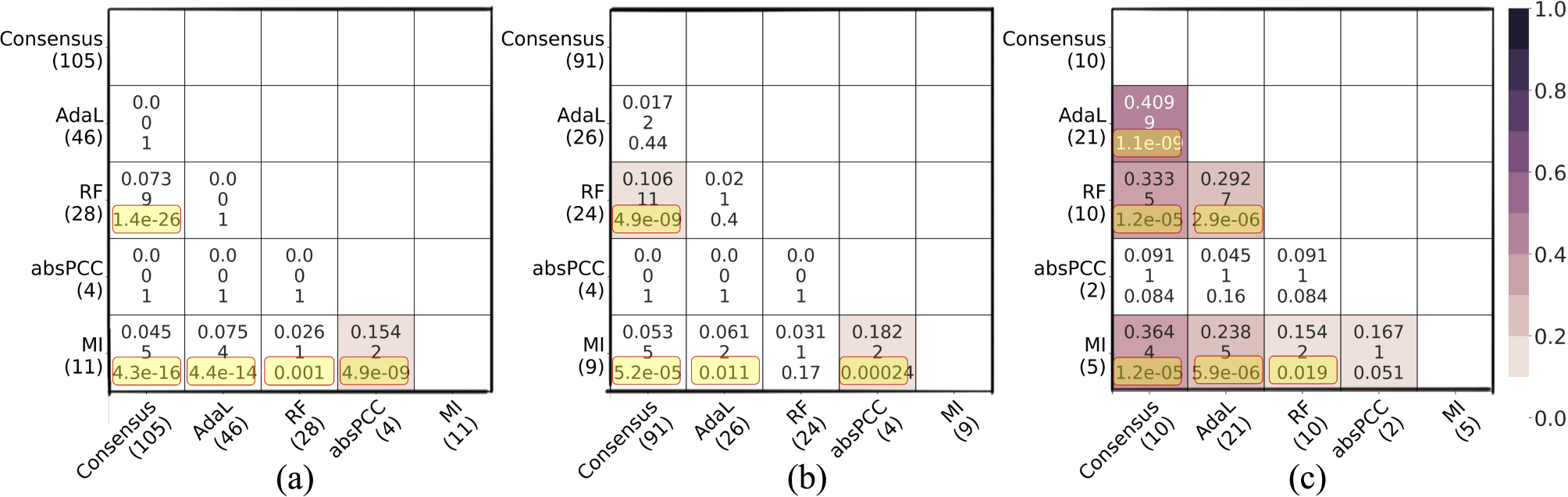
Pairwise overlaps between predictions made by the best combination for each of the five networks with respect to predicted novel associations, i.e., novel predictions. Panel (a) shows pairwise overlaps of gene-GO term associations. Panel (b) shows pairwise overlaps of unique genes that participate in the predicted gene-GO term associations. Panel (c) shows pairwise overlaps of unique GO terms that participate in the predicted gene-GO term associations. Within each panel, the number in parentheses below each network name is the number of corresponding novel predictions. Within each cell, there are three numbers; the first number is the Jaccard index, the second number is the raw number of the predictions in the given overlap, and the third number is the adjusted *p*-value resulting from the hypergeometric test. The yellow boxes highlight the adjusted *p*-values that are statistically significant.

For true positive predictions, in total, the five considered networks (i.e., combinations) predict 169 true positive gene-GO term associations, which involve 109 unique genes and 14 unique GO terms. While most of the pairwise overlaps are statistically significant (adjusted *p*-values ¡0.05), all Jaccard indices are low. That is, the lowest and highest Jaccard indices, respectively, are 3.3% and 27.2% for predicted true positive gene-GO term associations; 4.7% and 37.5% for unique genes that participate in the predicted associations; and 11.1% and 70% for unique GO terms that participate in the predicted associations (Fig. 4). For novel predictions, in total, the five networks predict 174 novel gene-GO term associations, which involve 131 unique genes and 26 unique GO terms. About half of the pairwise overlaps are statistically significant (adjusted *p*-values ¡ 0.05), but again, all Jaccard indices are low. That is, the lowest and highest Jaccard indices, respectively, are 0% and 15.4% for predicted novel gene-GO term associations; 0% and 18.2% for unique genes that participate in the predicted associations; and 4.5% and 40.9% for unique GO terms that participate in the predicted associations (Fig. 5).

The above results indicate that the predictions (with respect to both true positives and novel predictions) are largely complementary to each other. This observation strengthens our finding that the different network inference methods capture different biological signals. Importantly, despite the Consensus network not performing well compared to the individual networks in term of precision, it does uncover gene-GO term annotations not found by the other networks.

We conclude this section by qualitatively complementing the quantitative results thus far on the overlap between gene-functional predictions of the different combinations, by breaking down the overlaps by biological processes. Here, for comprehensiveness, we go back to all possible combinations. Namely, recall that we deal with the 11 constructed networks, two clustering methods, and up to three clustering parameter values. For each of the 11 *×* 2 = 22 combinations of network and clustering method, we consider any gene-GO term association predicted by at least one of the three clustering parameters. Then, we take the union of all such predictions over all combinations. Of all of the predictions in this union, we focus on those that are in the ground truth data, i.e., we consider the true positives over all combinations. We do this because true positives can be trusted more than novel predictions. Finally, we measure, for each of the 22 combinations of network and clustering method, the percentage of all of the true positives from the union that a given combination predicts.

We show these results in Fig. 6, which are broken down by individual GO terms that are then grouped into biological process categories based on the GO terms’ semantic similarities (see Section *“Methods”* for details). This analysis is intended to complement results from Fig. 4(a) on how much true positive predictions of different combinations overlap, by also providing insights into what biological process(es) the overlaps come from. From Fig. 6, we find that the overlaps between the different combinations of network and clustering method correspond mostly to pathogenesis-related GO terms, as well as to some of the GO terms related to cell cycle and transcription/translation. However, for most of the GO terms *not* related to pathogenesis, the different combinations of network and clustering method yield at least somewhat complementary results. The fact that most of the combinations capture the pathogenesis-related GO terms well is encouraging, as these biological processes annotate genes known to be involved in *P. falciparum* infection and immune response.

**Figure 6.**
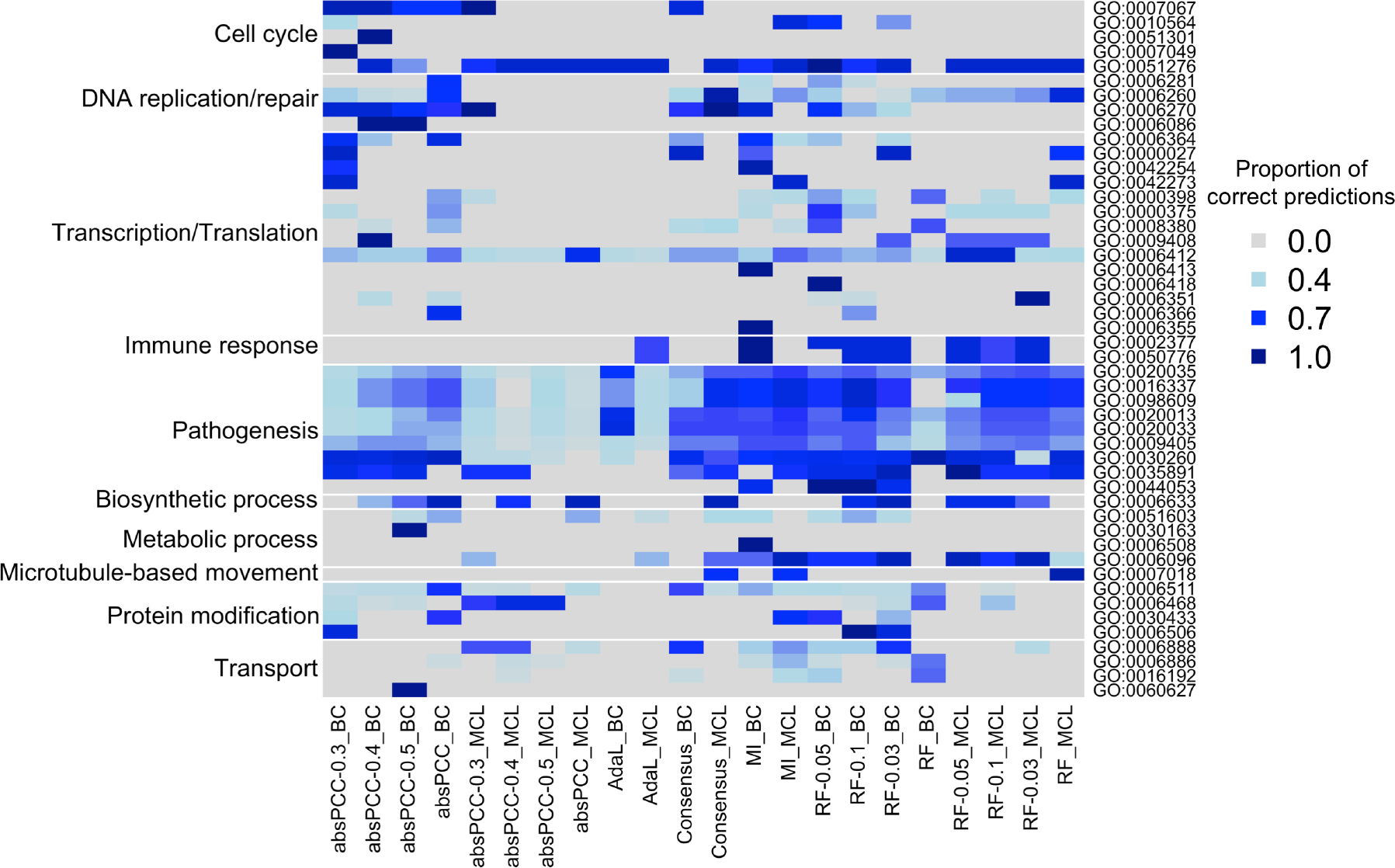
GO terms and biological process categories captured by the different combinations of network and clustering method. Columns correspond to the different combinations. Rows correspond to GO terms that are captured by predicted gene-GO term associations, where the GO terms are then grouped into biological process categories (shown on the left). We visualize the number of gene-GO term associations predicted by a given combination divided by the total number of true positives predicted over all combinations. That is, each cell shows, for a given GO term, the proportion of true positives from the union of all combinations that are predicted by a given combination. The darker blues represent higher proportion values.

### Ranking predicted gene-GO term associations and gene-gene interactions

Next, we aim to supplement the limited functional annotation data in the *P. falciparum* genome. Again, here, we deal with the 22 combinations of network and clustering method, with up to three clustering parameter values for each combination. For each of the 22 combinations, for each gene-GO term association predicted by a given combination, given up to three three adjusted *p*-values for a given prediction (corresponding to up to three clustering parameter values), we select the lowest adjusted *p*-value for the prediction. Then, we assign a confidence score to each gene-GO term prediction made by at least one of the 22 combinations of network and clustering method and provide the list of all associations ranked by their confidence scores. Intuitively, the more combinations support a predicted gene-GO term association and the more strongly a given combination supports a prediction (i.e., the lower the corresponding selected adjusted *p*-value), the higher the confidence score of the prediction (see “Methods”). We provide two ranked gene-GO term associations lists, one for existing associations with 1,062 such associations (Additional File 2) and the other for novel associations with 28,826 such associations (Additional File 3). For the distribution of confidence scores, see Fig. 7.

**Figure 7.**
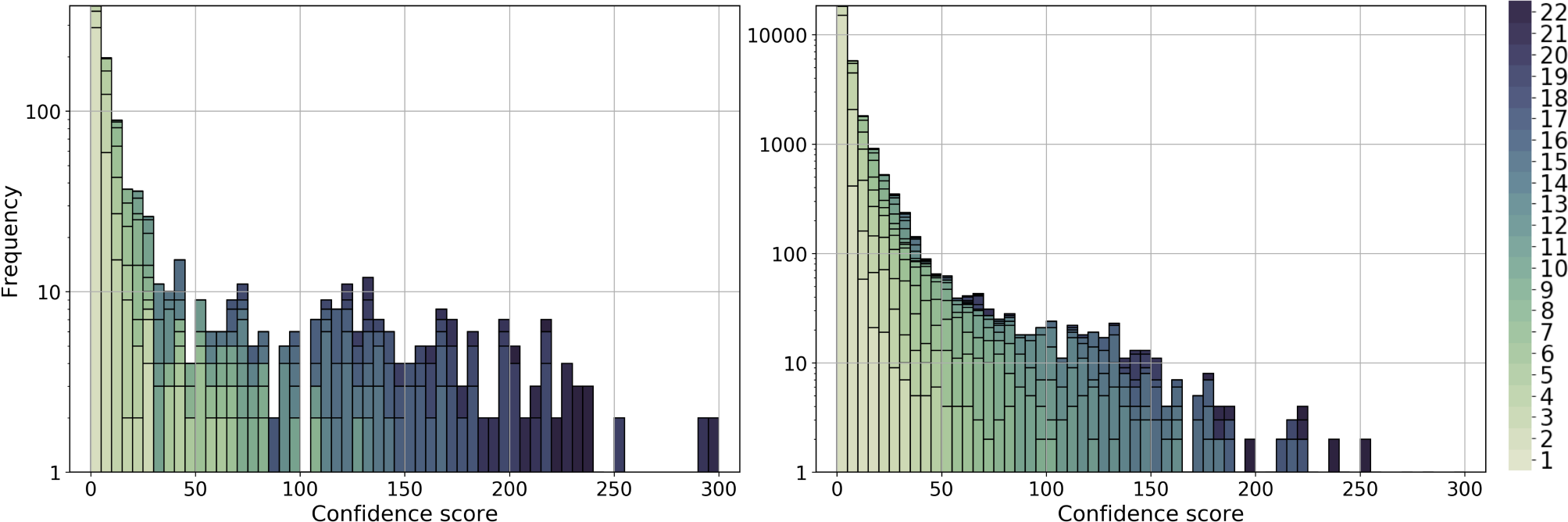
The distribution of confidence scores for predicted gene-GO term associations. The left panel shows the distribution for existing associations, i.e., true positives, and the right panel shows the confidence score distribution for novel associations. The color shades represent the number of combinations of network and clustering method that support the corresponding association. The darker color the color, the higher the support. Analogous results for gene-gene interactions are shown in Supplementary Fig. S8.

Similarly, to supplement the limited gene-gene interactions data in the *P. falciparum* genome, we provide a ranked list of gene-gene interactions with their confidence scores (Additional File 4). Intuitively, given all statistically significantly enriched clusters from the 22 combinations of network and clustering method, we consider all genes from the clusters along with their edges from the corresponding network. Then we assign to each edge a confidence score; intuitively, the more clusters contain a given edge (i.e., one of its end nodes), and the more functionally meaningful a given cluster is (i.e., the more GO terms it is enriched in and the lower the corresponding adjusted *p*-values of the enrichments), the higher the confidence score of the edge (see *“Methods”*). The ranked list includes 1,018,420 gene-gene interactions. We visualize the distribution of the interaction confidence scores in Supplementary Fig. S8.

### Validating the inferred networks using Endocytosis-related biological signatures

Endocytosis is an essential biological process for trafficking extracellular material in the cell to specific organelles. Extracellular material is first brought into the cell in early endosomes that mature into late endosomes as they are directed to lysosomes or through trafficking pathways from the Golgi apparatus. K13 and EPS15 localize to the periphery of the cell and are suggested to contribute to the generation of early endosomes. Clathrin has been shown to function in *Plasmodium* and other apicomplexa in an atypical role, primarily through trafficking pathways of the Golgi apparatus [20, 54]. A recent study [7] identified genes that interact via PPIs with each of K13, ESP15 and clathrin. The K13 and EPS15 gene lists were devoid of clathrin, suggesting that K13 functions in a clathrin-independent endocytosis pathway. Our co-expression networks may provide a systems-level perspective on these recently published data, including new hypotheses for further exploration into this cellular pathway and the mechanism of resistance for artemisinin.

For each of the K13, EPS15, and clathrin gene lists, we measure how densely the genes in a given list are connected to each other as well as to the genes in the other lists in the Consensus network. Note that even though the Consensus network does not perform the best in terms of precision in the task of predicting gene-GO term annotations, it does cover complementary edges from the individual networks. As such, here we aim to validate the effectiveness of the Consensus network from the endocytosis perspective. We measure the density of connections between gene members of each list, between genes that are in the union of each pair of lists, and between genes that are in the union or all three lists. Note that if a gene belongs to more than one of the K13, EPS15, and clathrin lists, we do not consider such a gene in this analysis.

We find that the genes in the K13 list are the most densely connected to each other, followed by the genes in the union of the K13 and EPS15 lists (K13-EPS15), followed by the genes in the union of the K13 and clathrin lists (K13-clathrin), and finally followed by the genes in the union of all three lists (All-Endocytosis) (Fig. 8). Genes within the K13 list are expected to be densely connected to each other in the network, and so are genes in the K13-EPS15 set, because the K13 and EPS15 gene lists together, while at the same time being devoid of clathrin, contribute to the generation of early endosomes [7]. This means that the Consensus network identifies expected pathways and their interacting proteins, which validates the network. The dense interconnectivity between the genes in the K13-clathrin set is a more interesting result because K13 does not directly interact with clathrin, and it would not necessarily be expected that the K13- and clathrin-interacting genes share the same network. The majority of the material imported by the K13-defined endocytosis pathway is host cell hemoglobin, which is eventually transported to the food vacuole, a lysosome-like structure where hemoglobin is degraded. It is plausible to hypothesize that endosomes generated by K13 are integrated into the trafficking network coordinated by clathrin and its interacting proteins through the Golgi, a result that would suggest the separate gene functions in related biological pathways even though they do not directly interact with each other.

**Figure 8.**
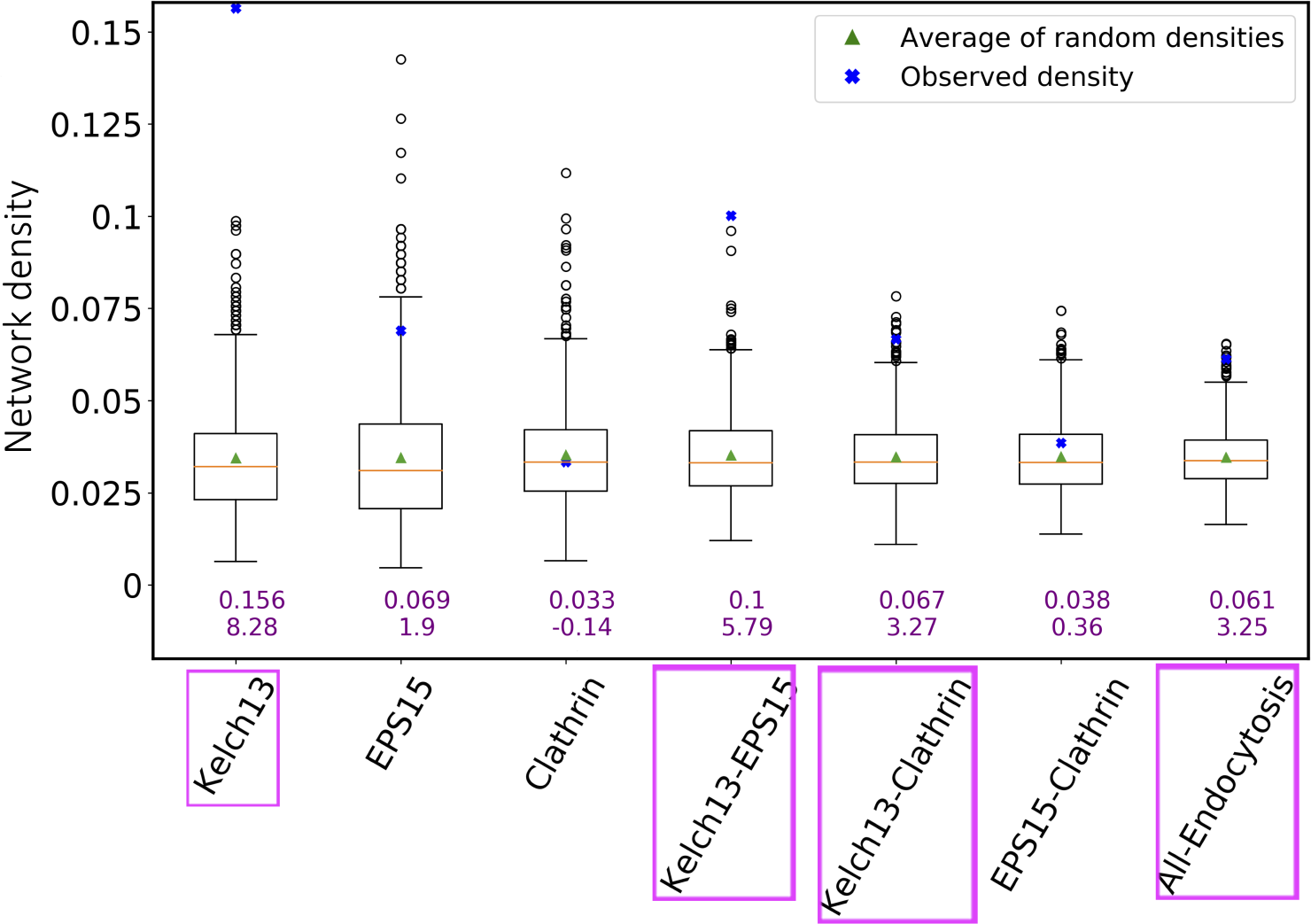
The connectivity of genes in a given list/set (or simply group) compared to the genes’ connectivity expected by chance. The *x*-axis shows the considered endocytosis-related gene groups. The *y*-axis corresponds to connectivity as measured by network density (for a gene group of size *n*, where there exist *e* edges between the *n* genes, the density is the ratio of *e* and 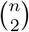, where the latter is the total possible number of edges between *n* nodes). The boxplot for a given gene group represents the density distribution of 1,000 random runs for that group. In particular, if a given gene group has *n* genes, the connectivity by chance is measured by calculating the density of a subnetwork consisting of randomly selected *n* genes and their edges from the Consensus network. The green triangle in each boxplot/for each group is the average density of the 1000 randomly selected subgraphs. The orange line is the median of the 1,000 random densities. The blue cross point is the actual (observed) density of the given gene group, whose numerical value is also shown as the top purple number below the given boxplot. The other (bottom) purple number below the given boxplot is the *z*-score of the observed density when contrasted against the random densities. The names of endocytosis-related gene groups that are shown in pink ovals correspond to those groups that are significantly more densely connected than at random).

Furthermore, the results suggest the three gene lists have potential to be operating together in the endolysosomal system to bring material into the cell and transport the material to its destination farther in the cell. Our analysis provides deep, testable extensions of the observed PPI data from [7] for how K13, EPS15 and clathrin are functioning as a system beyond what can be learned from only considering directly-interacting protein partners.

Interestingly, the genes in the EPS15 list alone, in the clathrin list alone, or in the two lists combined are not statistically significantly interconnected to each other when analyzed separately from the K13 gene list. This fits a scenario of EPS15 and clathrin having pleiotropic functions and operating in multiple pathways. It is likely that the global transcript data used for the networks does not capture the broad spectrum of interacting partners that either protein has throughout the complex *P. falciparum* cell cycle. Analyzing the gene lists together provides greater context to specific functions in the cell cycle when K13 is essential for cell development. Co-expression networks for specific stages of the *Plasmodium* cell cycle could further define the multiple functions that EPS15 and clathrin preform in the cell. Endocytic mechanisms in *P*. falciparum have not been fully elucidated and are an important area of research [61]. The Birnbaum *et al*. [28] study is a significant step in understanding how K13 functions inside the cell and it suggests that K13 and clathrin perform different functions inside the cell in separate pathways. Our data suggests that the functional pathways for K13 and clathrin are more correlated than would be expected by only the PPIs found in Birnbaum *et al*. [28]. Further molecular experiments will provide greater context for how the two functional pathways are coordinated and influence artemisinin resistance.

## Conclusions

In this study, we have constructed gene co-expression networks for *P. falciparum* using four prominent network inference methods. Then, we have evaluated the inferred co-expression networks in terms of their ability to predict existing functional knowledge (i.e., gene-GO term associations) through network clustering and leave-one-out cross-validation. Our results show that the different networks capture complementary gene-gene co-expression relationships (i.e., interactions) and also predict complementary gene-GO term associations. We have provided ranked lists of inferred gene-gene interactions and predicted gene-GO term annotations for potential future use by the malaria community. Our study indicates that there does not exist a gold-standard co-expression network which captures all aspects of the *P. falciparum* biology. Thus, relying on a single network inference method should be avoided when possible. In fact, we have demonstrated that the Consensus network which combines the interactions from the complementary individual co-expression networks agrees with the biology of the endocytosis-related cellular pathways and could thus yield new hypotheses about the mechanism of resistance for artemisinin.

## Methods

### Data

#### Gene expression data

We use gene expression data (GSE19468) to construct our co-expression networks curated by Hu *et al*. [21] in 2009, as follows. Drug perturbations were conducted for 29 drugs across 10 experimental groups (each with a no drug control and two - four drugs), see Table 3). Samples were collected at five - ten time points across the *P. falciparum* intra-erythrocytic developmental cycle. This experiment resulted in *P. falciparum* transcription profiles for 247 samples. Transcript abundance levels were obtained using a spotted oligonucleotide microarray [22] with 10,416 probes representing the 5,363 genes in the PlasmoDB *P. falciparum* genome version 4.4. In this unprocessed dataset probes are in rows and samples are in columns. There have been major revisions in subsequent PlasmoDB *P. falciparum* genome versions. So, rather than using the processed transcription profiles directly, we reprocessed the probe level data to better reflect genome updates and improved normalization methods.

**Table 3.** The information about drug treatments and time courses of GSE19468. For example, group 1 has eight time courses for one control group and three drug treatments. That is, there are 8 *×* 4 = 32 gene expression levels for a given gene obtained from group 1. By combining the expression levels across all groups, in total, there are 8 *×* 4 + 5 *×*4 + 7 *×* 5 + … + 10 *×* 3 = 247 gene expression levels for each of the 5,075 genes. Of which, 8 + 5 + 7 + … + 10 = 64 gene expression levels are from control treatments.

| Group | Time Points | Treatments |
| --- | --- | --- |
| 1 | 8 | Control, RoscovitineA, CyclosporineA, FK506 |
| 2 | 5 | Control, Colchicine, Na3VO4, StaurosporineA |
| 3 | 7 | Control, ML7, W7, KN93, Staurosporine |
| 4 | 6 | Control, Artemisinin, Chloroquine, Febrifugine, Quinine |
| 5 | 5 | Control, E64, Leupeptine, PMSF, RetinolA |
| 6 | 6 | Control, Apicidin(troph 5nM), Apicidine(troph IC90) |
| 7 | 5 | Control, Apicidin(schiz IC50), Apicidin(schiz IC90) |
| 8 | 6 | Control, TrichostatinA(IC50), TrichostatinA(IC90) |
| 9 | 6 | Control, Chloroquine(IC50), Chloroquine(IC90), Chloroquine(2*IC90) |
| 10 | 10 | Control, EGTA(IC50), EGTA(IC90) |

#### Processing probe-level data

To reflect revisions to the *P. falciparum* genome since 2009, nucleotide sequences for the 10,416 probes on the spotted array were obtained from GEO (GPL7493) and were aligned to the PlasmoDB *P. falciparum* genome version 36 using blast+ (v2.6.0) from NCBI. The 9,870 probes that aligned the *P. falciparum* transcriptome (PlasmoDB v36) with perfect match (bitscore ¿= 130) and no secondary alignments (secondary bitscores all ¡ 60) were retained in the dataset. These probes aligned to transcripts for 5,075 genes. Probes aligned to transcripts for the same gene were averaged and genes with non-zero values in ¿ 80% of samples were retained in the dataset. This data processing introduced missing values into the dataset. After blast mapping, we end up with 4,502 genes with gene names updated in our gene expression data (Additional File 5 - GSE19468_blasted.csv).

#### Imputing missing values in the gene expression data

Some of the network inference methods utilized in this study cannot be used when the underlying gene expression data contain missing data e.g., absPCC, thus it was necessary to impute these missing values. We test seven prominent imputation methods designed for gene expression data. Then, we select the method that performs the best in the expression data used in this study. By “best”, we mean the method that yields the smallest normalized root mean squared error (NRMSE). In particular, 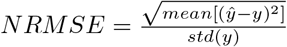, where *ŷ* represents the imputed values and *y* represents the actual values. The seven prominent imputation methods are explained as follows.

- Multiple imputations by chained equations (<u>MICE</u>) [8] imputes a column (i.e., sample) by modeling each sample with missing values as a function of other samples in a round-robin fashion. That is, given a sample column of interest, namely, *y*, and all other sample columns, namely *X*. A regressor is then fitted on *X* and *y* by learning a regression model from known values in *X* and *y* to predict the missing values in *y*.
- <u>SVDimpute</u> [69] is a singular vector decomposition (SVD)-based imputation method. Intuitively, a matrix can be recovered asymptotically by only using the significant eigenvalues. That is, given a gene *u*, a regression model of gene *u* and *k* most significant eigenvalues (i.e., eigengenes) are fitted. Then, the learned coefficients of the linear combination of the *k* eigengenes are used to impute the missing values of gene *u*. The processes are repeated iteratively until all missing values were imputed.
- <u>KNNimpute</u> [69] imputes missing values as follows. First, given a gene *u* with a missing value in sample *j, k* other genes without a missing value in sample *j* and are most similar to gene *u* are selected. Then the weighted average expression level of the *k* selected genes in sample *j* is treated as an estimated expression level for gene *u* in sample *j*, where the weight is the expression similarity (measured using Euclidean distance) of a gene (i.e., among the *k* selected genes) to the gene *u*. We vary *k* from 1 to 24 with an increment of 2.
- Local least squares imputation (<u>LLSimpute</u>) [29] imputes missing values as follows. First, *k* other genes that are most similar to (i.e., have the largest absolute Pearson correlation coefficients with) gene *u* of interest are selected. It differs from KNNimpute (i.e., *k* is predefined) that the *k* value for LLSimpute is introduced automatically.
- <u>SoftImpute</u> [45] imputes missing values by guessing values repeatedly. Specifically, the missing values in the gene expression data were initially filled as zero. Then a guessed matrix is updated repeatedly by using the soft-threshold SVD with different regularization parameters. If the smallest of the guessed singular values is less than the regularization parameter, then the desired guess is obtained. Please refer to [45] for methodological detail.
- <u>BiScaler</u> [19] was proposed based on SoftImpute but using alternating minimization algorithms. It introduced the quadratic regularization to shrink higher-order components more than the lower-order components such that it offers a better convergence compared to SoftImpute. Please refer to [19] for methodological detail.
- <u>NuclearNormMinimization</u> [10] imputes missing values by solving a simple convex optimization problem. That is, for a matrix *M* based on a theory that the missing values can be recovered if the number of missing values *m* obeys *m ≥ cN* ^1.2^*rlogN*, where *N* is the number of rows in matrix *M*, *c* is a positive numerical constant, and *r* is the rank of *M* . This algorithm usually works well on smaller matrices.

We use the library “pcaMethods” from Bioconductor R package [62] to perform LLSimpute and the python library “fancyimpute” to perform the remaining six imputation methods. We evaluate the performance of each method on our data as follows. (i) We take rows and columns without any missing values from the expression data as our ground truth data. (ii) We randomly remove 3.14% values from the ground truth data. We use 3.14% because it is the percentage of missing values in our expression data. We repeat this process five times and obtain five testing gene expression data. (iii) We apply an imputation method to each of the five testing data and compare the imputed matrix with the ground truth matrix. (iv) We select the method that yields the smallest average NRMSE across five runs. (v) Finally, we use the selected “best” method to impute the missing values for our entire gene expression data. It turns out that the MICE (with NRMSE = 0.204) is the best imputation method for our expression data.

#### Accounting for cyclical stage variation

*P. falciparum* has strong cyclical patterns of transcript expression across the intraerythrocytic development cycle (IDC) and these changes in gene expression tend to swamp other sources of transcriptional variation. To control for this strong cyclical variation we normalize each drug expression time point by its matched control time point. Since these are already log2 normalized expression profiles we subtract the control treatment from the experimental treatments for each time point. Specifically, for a given gene when treated by a drug, the *updated* expression level at time point *t* is obtained by using its *original* expression level minus the expression level at its control treatment at time point *t*. The resulting processed gene expression data thus has 4,374 genes and 183 (i.e., 247 *−* 64) expression levels (i.e., no control groups as these are used to normalize for cyclical variation). The corresponding data is attached as (Additional File 6 - GSE19468_final.csv).

#### Ground truth GO term annotation data

We use GO term annotations as our ground truth data to evaluate whether a cluster is statistically significantly enriched in a GO term. GO terms describe the knowledge of the biological domain with respect to three aspects: 1) molecular function, 2) cellular component, and 3) biological process. We focus on biological process because these terms group genes related to a single objective and is closest to defining gene products involved in the same pathways. We obtain all GO terms that describe biological processes and their annotated genes from two most commonly used databases for *P. falciparum* genes, i.e., GeneDB^3^ and PlasmoDB^4^. Because these two databases cover gene-GO term annotations that are complementary to each other, we combine their annotations and remove duplicates. As such, we obtain 4,736 gene-GO term associations that encompass 793 unique GO terms and 2,624 unique genes.

We use the processed data of gene-GO term associations to validate our clusters. Specifically, we treat each GO term as a ground truth cluster such that all genes annotated by this GO term belong to the same cluster. In general, a valid cluster should include at least three genes. So, we further process the ground truth data by only keeping those GO terms that annotate at least three genes in the expression data. We denote such GO terms as relevant GO terms and those associations involved with relevant GO terms as relevant gene-GO term associations. In our experiments, if we mention GO terms, we mean relevant GO terms. We summarize the statistics of our ground truth data in Table 4. The corresponding data is attached as (Additional File 7 - gene-GO term associations.txt).

**Table 4.** Statistics of relevant GO terms and gene-GO term associations.

| # of genes | # of samples | # of relevant GO terms | # of relevant gene-GO term associations |
| --- | --- | --- | --- |
| 4,374 | 183 | 255 | 3,232 |

#### Endocytosis data

PPI data used for the investigation of endocytosis-related genes in the Consensus network was generated recently by Birnbaum *et al*. [7] in 2020. The study used a BioQ-ID method coupled with mass spectroscopy using asynchronous cultures to define PPI of three proteins either directly implicated in artemisinin resistance (K13) or as controls (EPS15 and clathrin) to fully investigate the molecular function of resistance genes. Initially the PPIs of K13 (i.e., the molecular marker for artemisinin resistance) were identified and a subset were confirmed by cellular localization by immunofluorescence. The *P. falciparum* homolog of EPS15 was identified and confirmed as an interacting partner of K13 [68]. As such, EPS15 was chosen as a positive control for the BioQ-ID experiment and the genes identified as the PPIs of EPS15 largely overlapped with genes of the K13 PPI list. This added further evidence *K13* was involved in endocytosis. The study further defined a PPI list for clathrin because it is the main structural protein for canonical endocytosis pathways. The genes of the PPI list for clathrin did not overlap with genes of the K13 or EPS15 PPI lists leading to the conclusion that K13 functions in an endocytosis pathway devoid of clathrin. Our analysis utilizes the 173 endocytosis-related genes from [7]. The K13 contains 63 proteins, 60 of which are present in our gene expression data. The EPS15 PPI list contains 49 proteins, 48 of which are in the gene expression data. The clathrin PPI list contains 61 proteins, 58 of which are present in our gene expression data.

### Inference of gene co-expression networks

Given the processed gene expression data (GSE19468), we apply four network inference methods, i.e., mutual information (MI) [44], absolute Pearson correlation coefficient (absPCC) [56], Random Forest (RF) [24], and adaptive Lasso (AdaL) [32]. Moreover, we construct a Consensus co-expression network by taking the union of networks inferred using these four methods. In this section, we explain how we infer these co-expression networks.

#### Network construction using ARACNe framework

We use ARACNe framework for three network inference methods: MI, absPCC, and RF. Specifically, MI quantifies the mutual dependence (either linear or non-linear) of two random variables, and absPCC quantifies the linear correlation between two random variables, see [56] for detail. RF [24] is a tree-based method aiming to recover a network involving *n* genes into *n* subproblems such that each subproblem recovers the regulation relationship of a given gene. We use the python program “GENIE3” to find the edge weight of each gene pair. For methodological detail, see the original publication [24].

Network construction processes of a network include weighting co-expression relationships for each node pair and finding an appropriate threshold to distinguish between edge and non edge. However, finding an appropriate weight threshold for real-world networks is challenging [56]. Different weight thresholds lead to different networks, hence different network topological structures [57]. Because our goal in this study is not to examine the effect of various weight thresholds but to uncover novel functional knowledge about *P. falciparum* using networks. So, we choose a well-established network construction framework called ARACNe [44] to construct three networks based on the aforementioned three edge weight strategies. ARACNe was originally proposed to infer gene regulatory networks, and has been successfully applied to infer gene co-expression networks [2, 3, 46, 60, 70]. ARACNe first calculates an appropriate threshold *I*0 based on an null hypothesis that gene pairs are independent to each other if their mutual information is below *I*0. Then network construction by ARACNe was conducted by first generating 100 bootstraps from the expression data. In each bootstrap, a certain number of microarray samples (i.e., in our case 183) for all genes are randomly selected with replacement and are permuted. Then, a network using mutual information is constructed and pruned using the pre-calculated threshold *I*0 and data processing inequality (DPI). In particular, given a triangle subgraph (i.e., genes *g*1, *g*2, and *g*3), DPI removes the edge with smallest weight in the triangle. Intuitively, this is because in this example, *g*1 and *g*3 are interacting with each other indirectly (i.e., through *g*2) and hence should be removed. Finally, the final network is obtained by keeping those edges that are detected across 100 bootstraps statistically significant amount of times. In other words, only non-random gene pairs are kept (i.e., adjusted *p*-values *<* 0.05 using false discovery rate (FDR) correction). For methodological detail, please refer to the original publication [44].

We directly apply the ARACNe-AP implemented by Lachmann *et al*. [33] to construct our MI co-expression network. For absPCC, we set the *I*0 to 0.6 according to [9, 27] that two metabolites with Pearson Correlation Coefficients *≥* 0.6 are considered as associated. Then we follow the ARACNe framework to obtain our final absPCC network using 100 bootstraps and the DPI technique. For RF, because the tree-based method includes a pruning step which functioned as a weight threshold, we do not set the threshold *I*0. Instead, we directly adopt the GENIE3 implementation from the original publication [24] and the DPI technique for each of the 100 bootstraps. Finally, we use the FDR correction from ARACNe framework to obtain the final RF co-expression network.

Because the resulting absPCC and RF networks are still very large, we further threshold these two networks by keeping *k*% most important edges out of all edges in the network without disconnecting the network (i.e., losing too many nodes in the largest connected component). We vary the *k* from 1-100 and select three such thresholds for each of the absPCC and RF networks. Thus, from this process, we obtain nine networks, i.e., four absPCC-based co-expression networks, four RF-based co-expression networks, MI co-expression network.

#### Network construction using Adaptive Lasso

Graphical Gaussian Models (GGMs) are prominent methods for modeling gene associations based on microarray gene expression data [12, 37, 59, 75]. GGMs are often used to obtain unbiased estimate of partial correlation between gene *i* and gene *j*, such that the partial correlation coefficient is treated as the edge weight of gene *i* and gene *j* in the resulting network. GGMs suffers two limitations, (i) they assume that the number of microarray experiments is much larger than the number of genes to ensure that the inversion of the covariance matrix in GGMs can be assessed statistically; (ii) they calculate partial correlation coefficients for all gene pairs in the expression data without a threshold. However, (i) the real-world gene expression data usually comes with a larger number of nodes compared to the number of microarray experiments; (ii) only those gene pairs with strong correlations (i.e., correlations coefficient greater than a certain threshold) are meaningful. To address such limitations of GGMs, Krämer *et al*. [32] proposed AdaL, a GGMs-based method that uses Lasso penalty to shrink the small edge weights (i.e., coefficients) to zero by using L1 norm regularization, i.e., Lasso. This way, those edges that are not meaningful will be shrunk to zero, i.e., removed from the resulting network. As such, those gene pairs with strong partial correlations are kept and hence form a network. For methodological detail of AdaL, please refer to the original publication [32]. Note that we do not apply ARACNe for AdaL as it already uses Lasso penalty to shrink small edge weights to zero. Thus, from this process, we construct another co-expression network, i.e., AdaL.

We use the R package “parcor” (available from the R repository CRAN) [32] to generate our AdaL network using our gene expression data. The input data is a *m × n*-dimensional matrix where *m* is the number of microarray treatments (i.e., 183) and *n* is the number of genes (i.e., 4,374). The output is a *n × n* adjacency matrix of the resulting network.

#### Construction of the Consensus network

Co-expression networks that are inferred using different inference methods have complementary edges (Section *“Results and discussion”*). So, we aim to examine the properties of a network that is the union of four aforementioned co-expression networks, namely, Consensus co-expression network. We construct the Consensus network using MI, smallest absPCC (absPCC-0.3), smallest RF (RF-0.03), and AdaL, as follows.

1. Given that we have four networks with different number of edges. If network 1 has *x*1 edges and network 2 has *x*2 edges where *x*1 *< x*2. We argue that the least important edge in network 1 (ranked as *x*1) should be equally important as the edges have rank *x*1 in network 2. This is because each network is inferred via methods with well-established thresholding strategy, and a network with more edges is intuitively have a more loosen thresholding strategy compared to a network with fewer edges.
2. Because we aim to make sure that the higher the rank value of an edge, the more important the edge is. For example, in network 1, an edge with rank *x*1 means the edge is the most important edge in network 1. So, we reverse the way we rank edges from step 1.
3. To assure the above two steps are satisfied when we construct the Consensus network, we first calculate the number of edges in all four networks (i.e., AdaL, MI, absPCC-0.3, and RF-0.03). If network 4 is the largest network with *x*4 edges, we use *x*4 as our possible maximum rank such that the most important edge in each of the four network has the same rank, which is *x*4 .
4. According to step 3, network 1 has edge ranks from *x*4 to *x*4 *− x*1, network 2 has edge ranks from *x*4 to *x*4 *− x*2, network 3 has edge ranks from *x*4 to *x*4 *− x*3, and network 4 has ranks from *x*4 to 1.
5. After we get all raw ranks for edges of each network, we use min-max normalization to normalize the rank of each edge in each network. That is, we first find the maximum (i.e., *max* also *x*4) and minimum (i.e., *min* also 1) edge rank across the four networks. Then, for a given edge between gene *i* and *j* with weight *wij*, the normalized rank is ((*wij − min*)*/*(*max − min*). The resulting normalized rank of edges across four networks spans from 0 to 1.
6. Finally, for a given edge, we sum the weights from the four networks. The collection of all such edges forms the Consensus network. That is, an edge between gene *i* and *j* in the Consensus network has a weight of *wij* = *sum*(*wijl*) for *l* = 1, 2, 3, 4, where *l* is the network id. Consequently, the resulting Consensus network has a maximum possible edge weight of 4 and a minimum possible edge weight of 0.

With the Consensus network, in total, we construct and analyze 11 co-expression networks in this study.

### Clustering methods

Recall that we test two prominent network clustering methods (i.e., cluster affiliation model for big networks (BigCLAM, referred to as BC) [78] and Markov Clustering (MCL) [14]) to predict gene-GO term associations. BigCLAM is a soft clustering method (i.e., a node can be assigned to multiple clusters) and MCL is a hard clustering method (i.e., a node can only be assigned to one cluster). We select one method for each type to reduce the effect of clustering type towards prediction accuracy of gene-GO term associations as much as possible. Also, we test multiple clustering parameters to give each network the best case of advantage.

#### BigCLAM

BigCLAM (BC) is a non-negative matrix factorization-based algorithm that assumes the overlaps between communities are densely connected. It also detects densely overlapping and hierarchically nested communities. For methodological detail and implementation, please refer the original publication [78]. We apply BigCLAM to each of the 11 networks inferred in this study. Specifically, for a given network, we run the BigCLAM with different parameters, i.e., the number of resulting clusters. We vary it from 50 to 700 with an increment of 25 from 50 to 500 and an increment of 50 from 500 to 700. Note that the number of resulting clusters is pre-defined number, meaning that the actual number of clusters can be different as the parameter. Then, we test the prediction performance of each network clustering parameter combination, and systematically select up to three combinations based on three pre-defined selection criteria, see *“Predicting and evaluating function annotations from clusters”* for detail.

#### MCL

MCL is an efficient random walk-based algorithm that assumes nodes that are densely connected to each other are similar to each other. MCL has been widely applied to detect the modules in the networks [5, 40, 76]. It takes the adjacency matrix representation and uses expansion and inflation stochastic process the make the densely connected area denser and the sparsely connected area sparser. We apply MCL to each of the 11 networks, and vary the inflation parameters within the range of 1.2 to 5.0 (i.e., as suggested from the user manual ^5^). A smaller inflation parameter tends to result in less number of clusters with bigger cluster sizes compared to a larger inflation parameter. Specifically, we test 33 inflation parameters with an increment of 0.02 for inflation from 1.2 to 1.4, an increment of 0.1 from 1.4 to 3, and increment of 0.2 from 3 to 4 and an increment of 1 from 4 to 5 for each network. Then we test the prediction performance each network clustering parameter combination, and systematically select up to three combinations based on three pre-defined selection criteria, see Section *“Predicting and evaluating function annotations from clusters”* for detail.

### Predicting and evaluating gene-GO term associations from clusters

We establish a systematic parameter selection and evaluation framework to evaluate each of the combinations of a network, a clustering method, and a clustering parameter value, as follows.

1. For the given gene expression data, we test against relevant GO terms to predict and evaluate the accuracy of predicted gene-GO term associations for each of the combinations.
2. For all clusters from a given combination, we use hypergeometric test (Section *“Hypergeometric test”*) to compute the probability scores (i.e., *p*-values) of the enrichment significance between each pair of a given cluster and a GO term. If a cluster is statistically significantly enriched in at least one GO term (i.e., *p*-value ¡ 0.05), we mark this cluster as an enriched cluster. We test all clusters and obtain the significantly enriched clusters.
3. In parallel, we make gene-GO term association predictions using significantly enriched clusters and GO terms via leave-one-out cross-validation [58], as follows.
  - First, we hide a gene *i*’s GO term knowledge at a time.
  - Second, we test whether each of the clusters that gene *i* belongs to is significantly enriched in any GO term. If such a cluster is statistically significantly enriched by a GO term *j*, we predict gene *i* annotated by GO term *j*.
  - Third, we repeat the above two steps for every gene that has at least one existing GO term annotation. Then we use precision, recall to evaluate prediction accuracy. The precision is the percentage of correct predictions among all predictions we make. The recall is the percentage of correct predictions among all existing gene-GO term associations. Because there is always a trade-off between precision and recall, we use precision as our parameter selection criteria (3). We do this because we believe that in biomedicine, for wet lab validation of predictions, it is more important to have a high precision if we can not have both high precision and recall [38, 39].
4. According to our three selection criteria, each combination has up to three clustering parameter values. Different selection criteria could end up with the same clustering parameter. This is why we have up to three selected clustering parameters for a given combination of a network and a clustering method.
5. Because we also aim to compare prediction performance across different networks. We first select the best clustering parameter based on their leave-one-out cross-validation precision and recall for each of the 11 networks. Specifically, for two clustering parameters, if parameter 1 has a higher precision and a higher recall or a similar recall compared to parameter 2, we select parameter 1. If parameter 1 has a higher precision but a lower recall compared to parameter 2, we keep both. For those selected “best” parameters of each network, we further compare them using the same selection criteria to find the combinations that yield the best prediction accuracy in terms of precision and recall.
6. We then qualitatively analyze our predicted gene-GO term associations using relevant biological pathways. In particular we visualize how effectively each of the 22 combinations predicts true gene-GO term associations as a heat map showing the proportion of gene-GO term associations correctly predicted for each GO term (rows) by a given combination (columns). GO terms were grouped using semantic similarity using a webtool called REVIGO [63]. Default REVIGO parameters were used to analyze the list of GO terms. The semantic similarity groupings and descriptions from REVIGO were exported by utilizing the “Export to TSV” option under the TREE MAP view.
7. Finally, we perform deep-dive analysis for those selected combinations from step 5 using Jaccard index and overlap coefficient. Both methods measure the overlaps between two sets. In particular, given set A and set B, the Jaccard index is 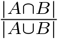, and the overlap coefficient is 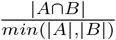 . The Jaccard index results in more accurate results when the sizes of set A and B are close to each other, while overlap coefficient results in more accurate result for small data when the sizes of set A and B are far away from each other. The number of predictions made by different combinations can be very different or similar, which is why we use both indices to quantitatively measure our deep-dive analysis.

#### Hypergeometric test

We perform hypergeometric test [17] for each pair of a cluster and a GO term (e.g., cluster 1 and GO term 1). Formally, the background (i.e., population) is the number of genes in the expression data of interest, represented as *N* . If *C* is a cluster, *G* is a GO term, *x* is the overlap between the cluster *C* and the GO term *G*, then the *p*-value is computed as: 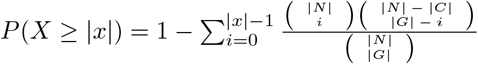. Because we perform multiple tests, we adjust the *p*-values using false discovery rate (FDR) correction as follows.

- Within each network cluster parameter combination, assuming we have *c*1 clusters and *g*1 GO terms. We adjust *c*1 *× g*1 *p*-values (i.e., each corresponds to a cluster and a GO term pair). We then use the adjusted *p*-values to determine whether a GO term is significantly enriched in a cluster.
- After we select up to six parameters per network based on our three systematic selection criteria, we recorrect up to 11*×* 6 *× c*1 *× g*1 *p*-values (i.e., we have 11 networks) for the purpose of fairly comparison between networks for a given gene expression data. By recorrect, we mean we take the up to 11 *×* 6 *× c*1 *× g*1 *p*-values and use FDR correction to obtain the adjusted *p*-values. We do not correct across all tested clustering parameters because some of the parameters are tested to make sure that we are not missing some of the important clustering parameters. Therefore, adding *p*-values from these parameters for test correction can be too conservative, and consequently remove lots of true positives.

### Assigning confidence scores to the predicted gene-GO term associations

In addition to evaluate the prediction accuracy of different networks, another main goal in this study is to predict and rank gene-GO term associations based on their confidence score. In particular, for a given gene expression data, we provide two lists of ranked gene-GO term associations, 1) existing association, i.e., true positive predictions (those that from ground truth data), 2) novel association predictions (those that do not currently exist), along with their “confidence” scores. Intuitively, the more of the networks support an association, and the more significant the association is (i.e., the lower the adjusted *p*-value of the association), the more confident the association is. We rank the associations as follows.

- Recall that we have 11 co-expression networks and two clustering methods, which totals to 22 combinations of a network and a clustering method (i.e., 22 adjusted *p*-values). Recall that we select up to three clustering parameters per combination, each corresponds to an adjusted *p*-value. We select the smallest adjusted *p*-value as the adjusted *p*-value for the corresponding combination.
- We rank predictions based on the number of combinations that support this prediction and their corresponding adjusted *p*-values. Specifically, we first take the negative log of the adjusted *p*-values (transformed *p*-values), such that the smaller the adjusted *p*-value a prediction has (i.e, the more important the prediction is), the larger the transformed *p*-value is. We then sum the 22 transformed *p*-values and get one final index for each prediction, i.e., confidence score. We rank the predictions from high to low based on their confidence scores.

### Assigning confidence scores to the predicted gene-gene interactions

Similar to the prediction of gene-GO term associations, we also provide a reference list of gene-gene interactions (GGIs) and their “confidence” scores. Intuitively, the interactions that yield the most meaningful gene-GO term associations, are supported by most amount of the networks, and/or are among the top-scoring interactions in their respective network, will be the most confident interactions. Specifically, the rank the GGIs as follows.

- For each of the 22 combinations, we first identify the statistically significantly enriched clusters, and get all genes in the clusters along with their edges from the network.
- For each identified edge, we assign the negative log transformed *p*-value associated with the cluster that the edge belongs to as its weight. If an edge belongs to multiple clusters, (e.g., genes can be grouped into multiple clusters via BigCLAM), we select the one with smallest adjusted *p*-value (i.e., the largest transformed *p*-value).
- We sum up the 22 transformed *p*-values (i.e., each correspond to a combination) and get a final confidence score for each GGI. We rank the GGIs from high to low based on their confidence scores.

### Examining the connectivity of endocytosis-related genes in the Consensus network

We use the Consensus network to examine the hypotheses of endocytosis process from Birnbaum *et al*. [7]. Specifically, if a network shows that the *K13* -related genes and *EPS15* -related genes are close to each other than at random in the network, it would corroborate the initial finding of the Birnbaum *et al*. study by a complementary analysis on independent data. We would also expect clathrin-related genes to be closer to each other than random and distant from *K13* -related and *EPS15* -related genes with connectivity similar to random in the network. We evaluate the connectivity of each group of Endocytosis genes using their network connectivity, i.e., the density, as follows.

- Given a group of Endocytosis genes, assume *m* genes, we select the induced subgraph (i.e., genes and their interactions) of the genes from a given network and we calculate the observed density of the subgraph. Network density measures how close a network is to its complete version (i.e., all pairs of nodes are connected).
- We then randomly select *m* genes from the network and their induced subgraph. We calculate the network density. We repeat this process a thousand times and calculate the z-score of the observed density compared to densities from the 1000 random runs.
- We use 2.0 as z-score threshold to determine whether the observed density is significantly larger than the random densities. In particular, if the z-score of a group of endocytosis genes is greater than 2.0, then the group of Endocytosis genes are densely connected than expected by chance.

## Supporting Information

**Supplementary Fig. S1.**
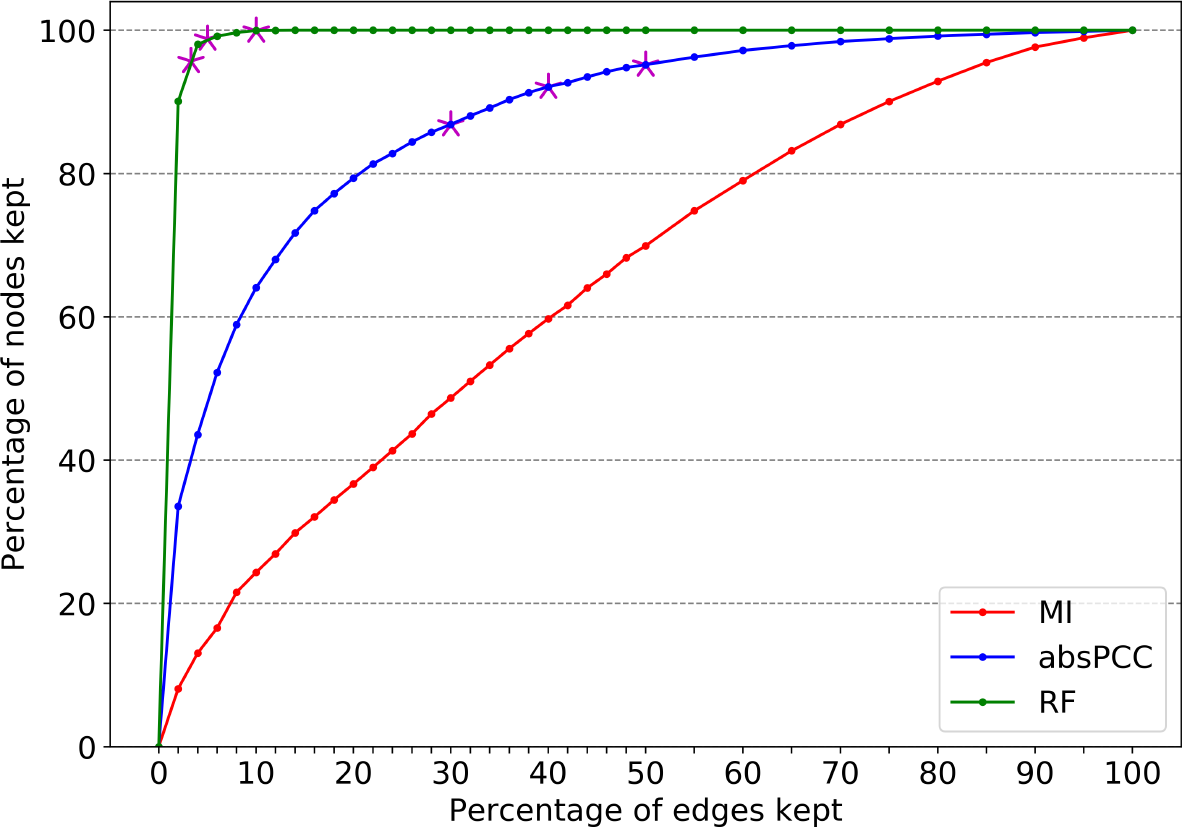
The effect of edge weight thresholds on the size of the largest connected component for a given network. It illustrates the percent of nodes remained in the largest connected component out of all nodes in a given network when keeping *k*% of highest weighted edges in the given network (i.e., MI, absPCC, RF). The x-axis represents the percent of highest weighted edges kept (i.e., *k*%) out of all edges in a given network. The y-axis represents the corresponding percent of nodes remained in the largest connected component out of all node in the gene expression data when keeping *k*% highest weighted edges. *k* is varied from 1 to 100. The magenta stars indicate the three thresholds we select for absPCC and RF, respectively. Note that we include MI in the figure as a control visualization for absPCC and RF.

**Supplementary Fig. S2.**
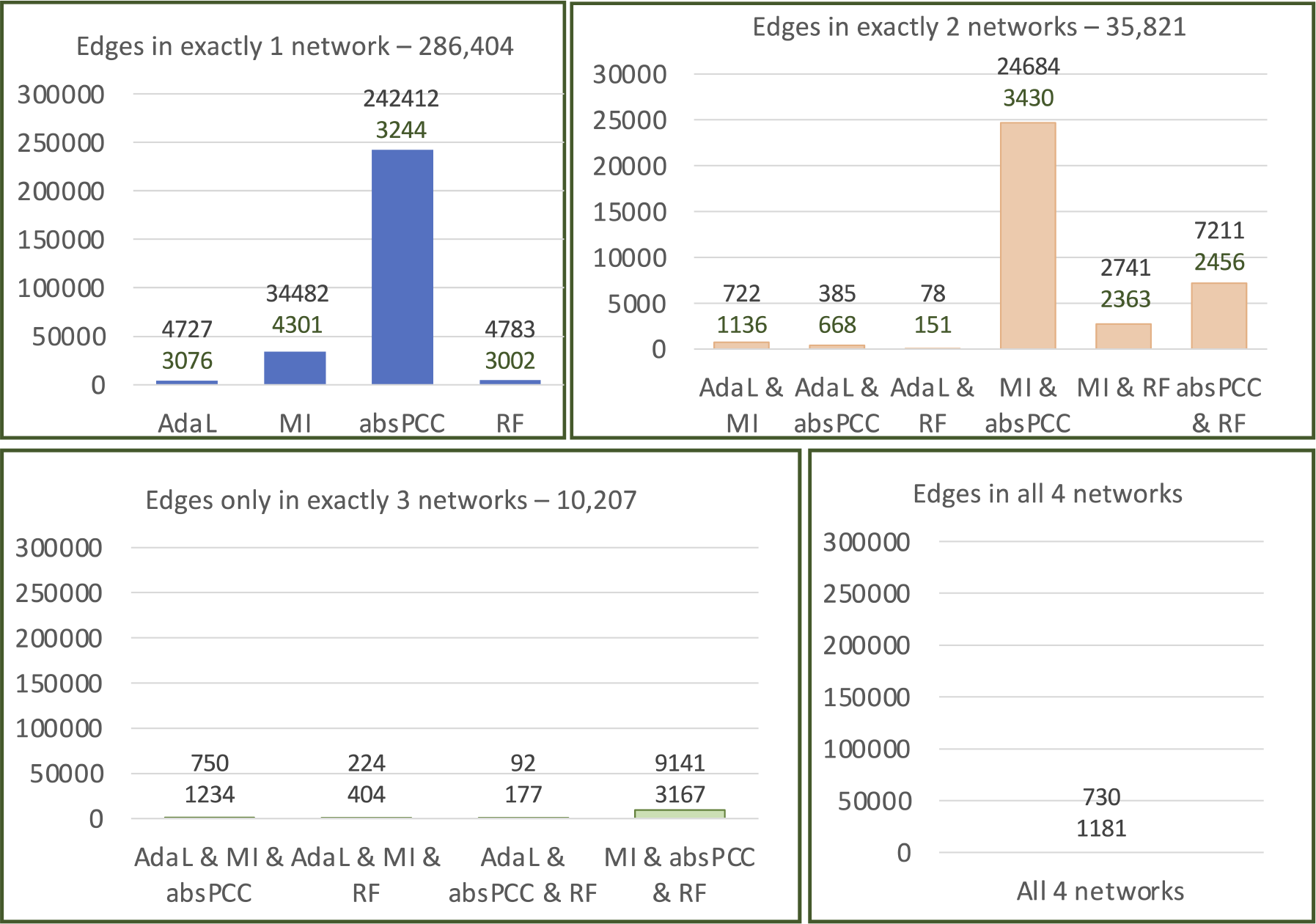
Details about pairwise edge overlap between the four networks, i.e., MI, absPCC, RF, and AdaL. The box on the top left is the number of unique edges that are only present in exactly one network, where the x-axis shows the networks. The box on the top right is the number of unique edges that are only present in exactly two networks, where the x-axis shows the network pairs. The box on the left bottom is the number of unique edges that are only present in exactly three networks, where the x-axis shows the network triplets. The box on the right bottom is the number of unique edges that are present in all four networks, where the x-axis shows the four networks. All y-axes show the edge counts. The two numbers on top of each bar across the four boxes represent the number of edges and the number of nodes these edges encompass.

**Supplementary Fig. S3.**
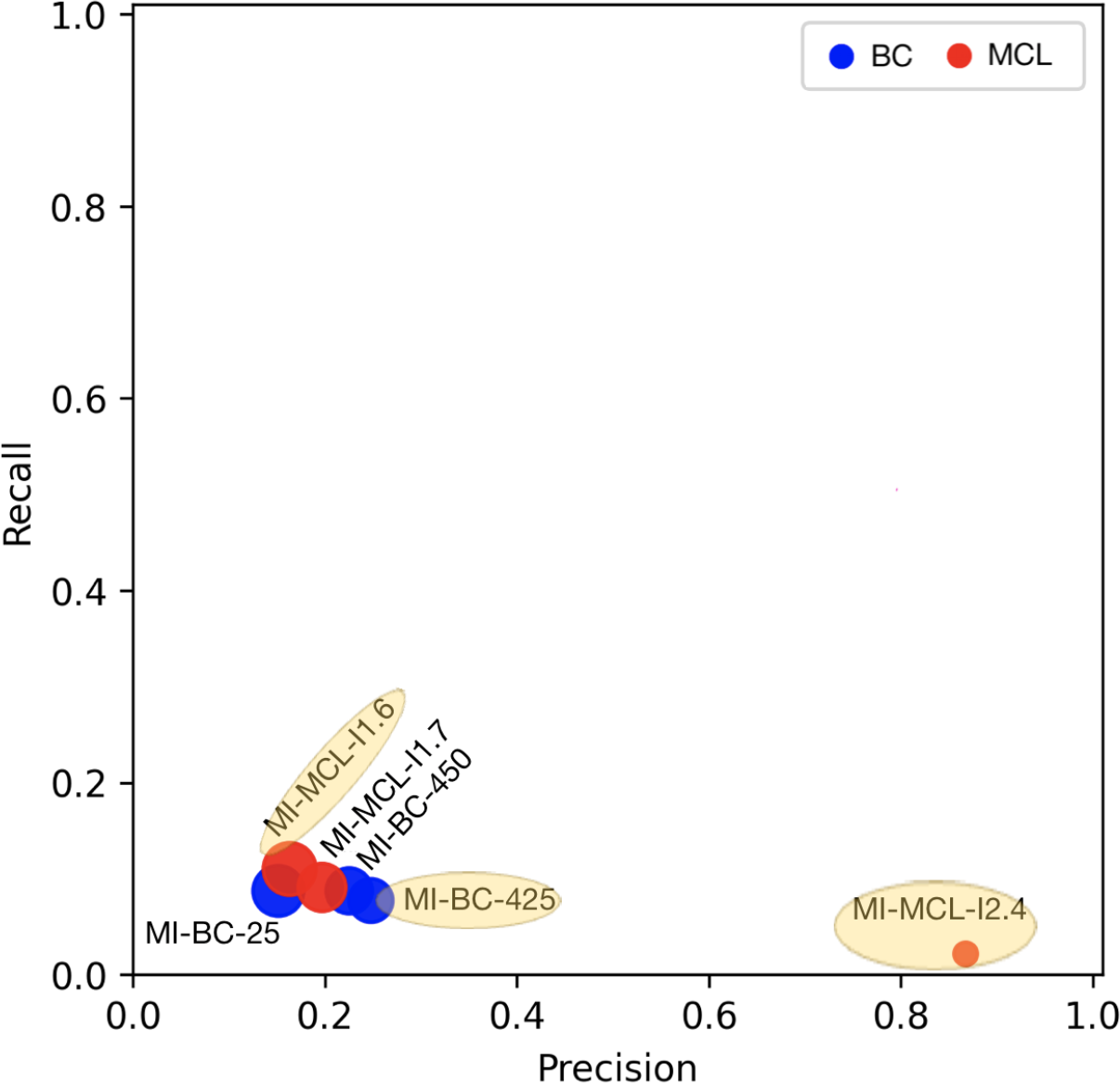
Accuracy of predicting gene-GO term associations in the leave-one-out cross-validation in terms of precision and recall via MI. The sizes of the points correspond to the numbers of predictions produced by a given combination. The color of the points corresponds to a clustering method.

**Supplementary Fig. S4.**
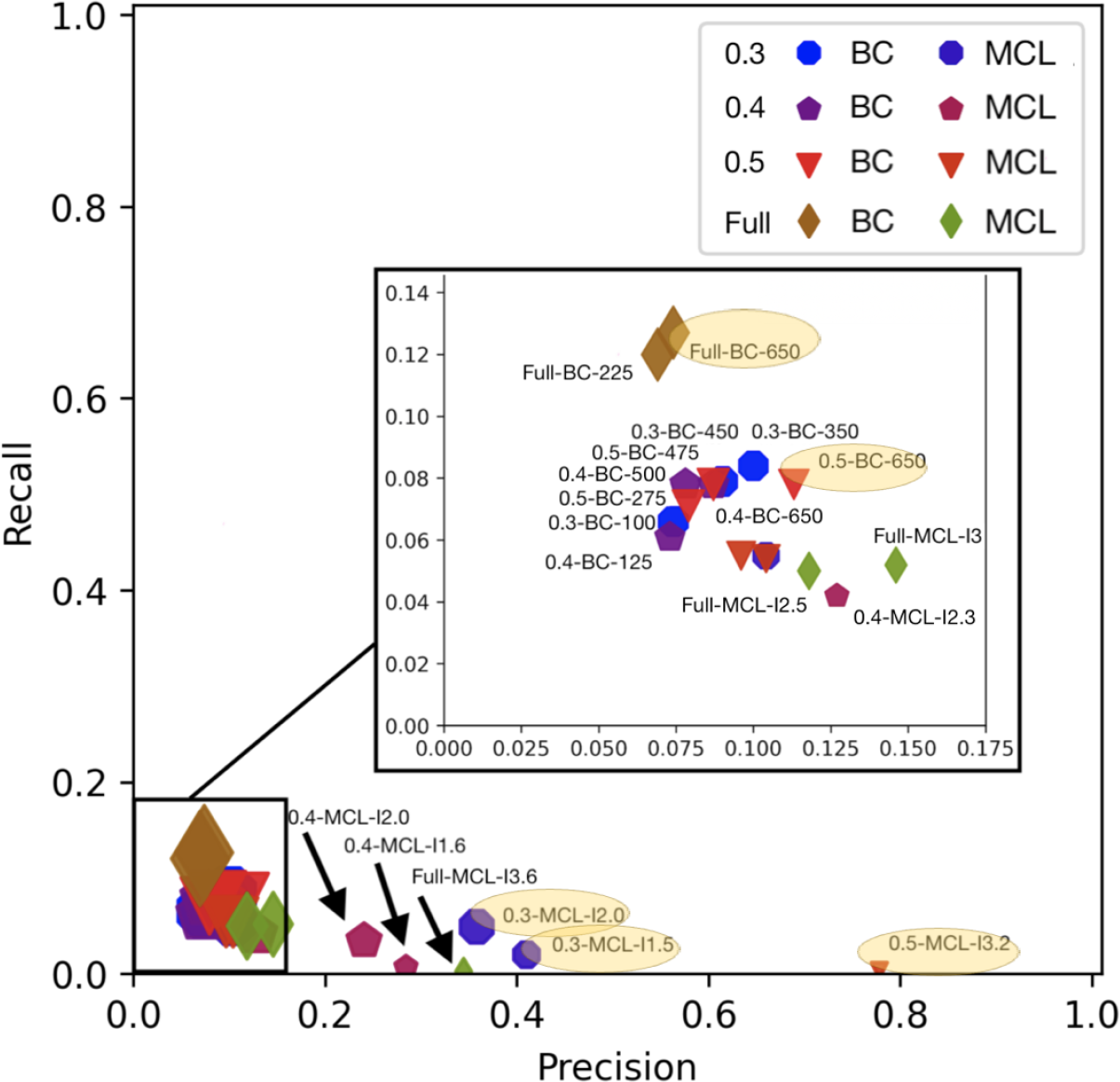
Accuracy of predicting gene-GO term associations in the leave-one-out cross-validation in terms of precision and recall via absPCC. Each point is a combination of network, clustering method, and parameter value. The sizes of the points correspond to the numbers of predictions produced by a given combination. The shape of a point corresponds to a network, and the color shade of a point corresponds to a clustering method. For example, all circles correspond to absPCC-0.3, of which, light blue corresponds to BC and dark blue corresponds to MCL.

**Supplementary Fig. S5.**
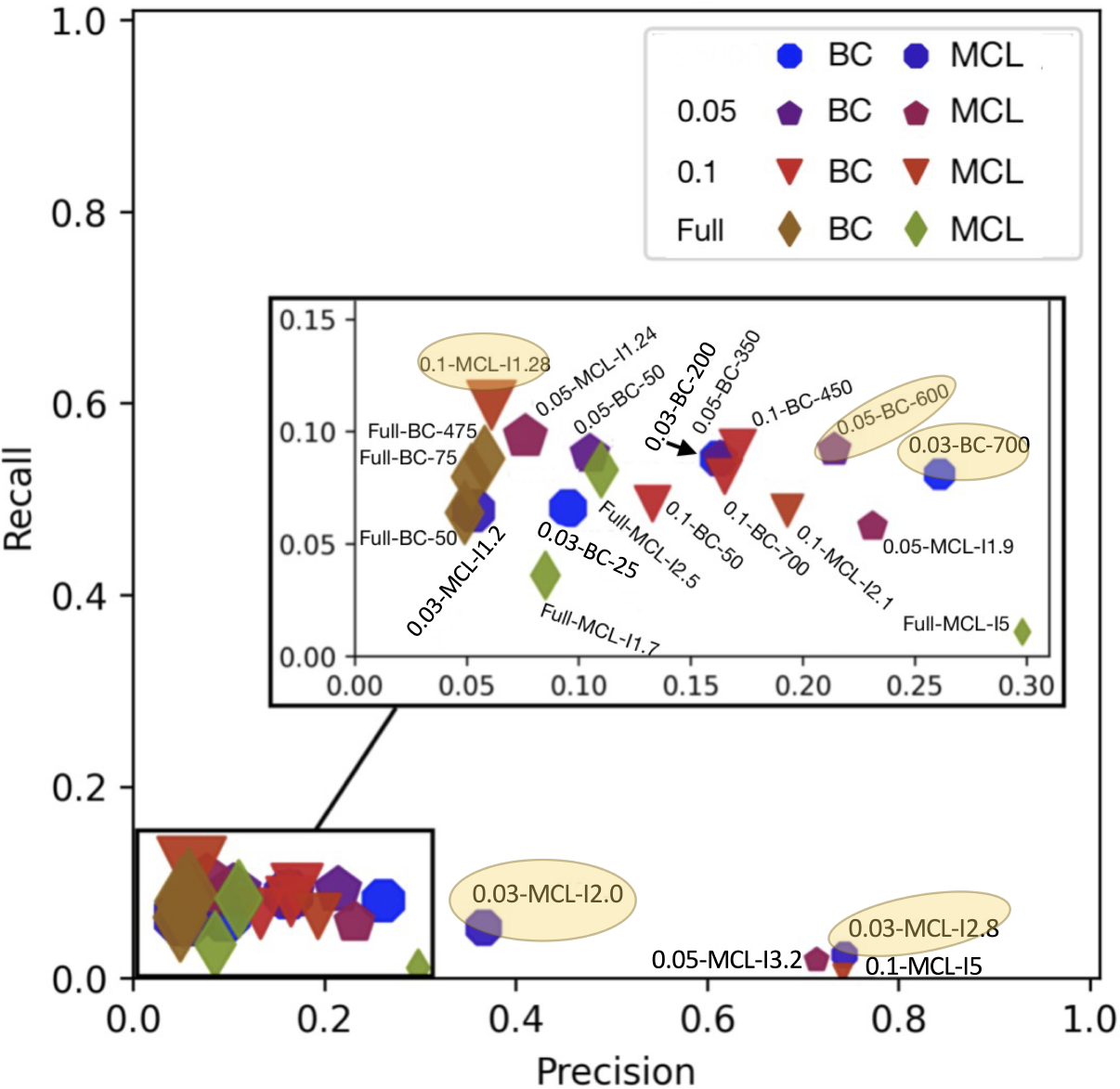
Accuracy of predicting gene-GO term associations in the leave-one-out cross-validation in terms of precision and recall via RF. Each point is a combination of network, clustering method, and parameter value. The sizes of the points correspond to the numbers of predictions produced by a given combination. The shape of a point corresponds to a network, and the color shade of a point corresponds to a clustering method. For example, all circles correspond to RF-0.03, of which, light blue corresponds to BC and dark blue corresponds to MCL.

**Supplementary Fig. S6.**
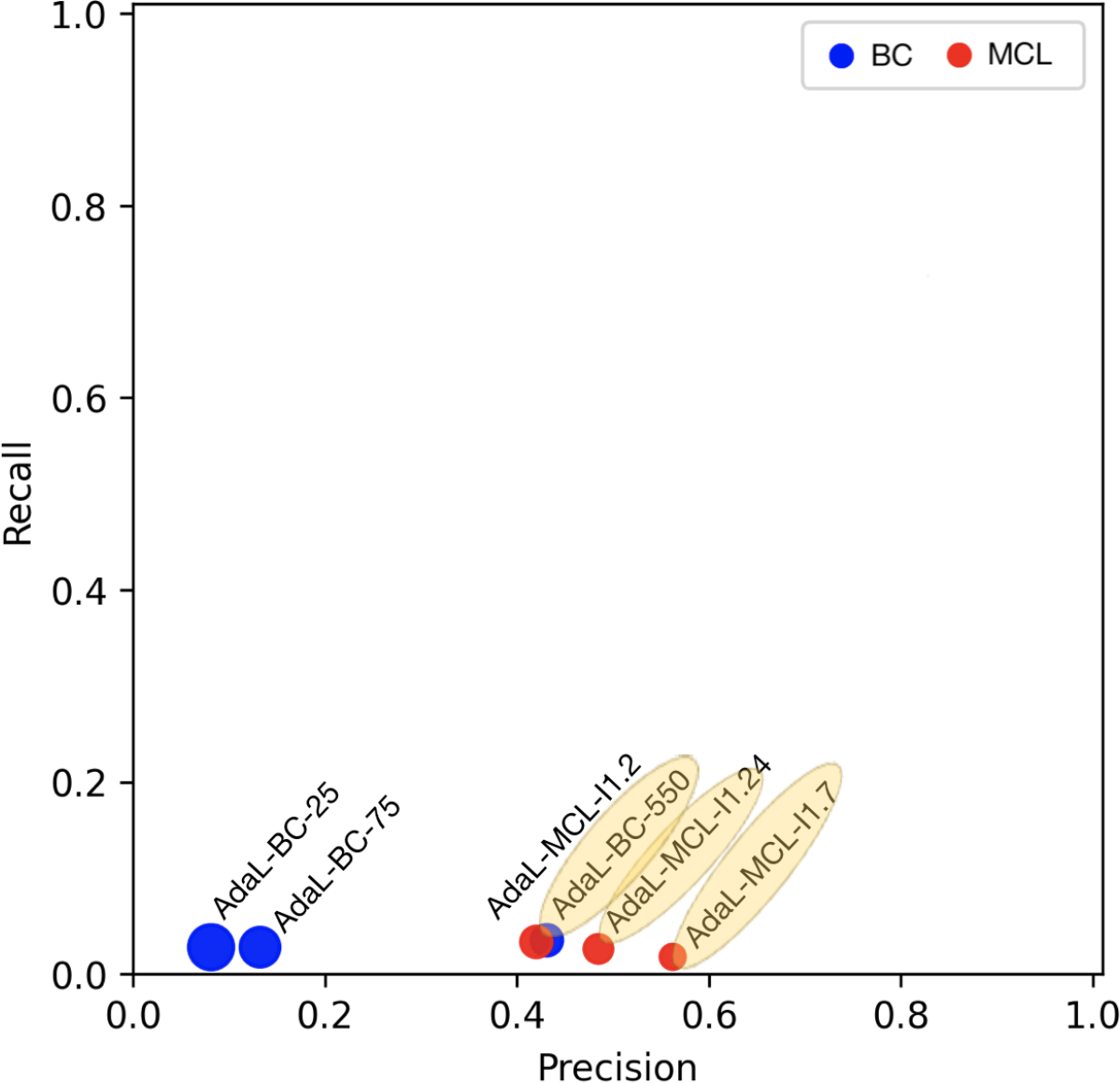
Accuracy of predicting gene-GO term associations in the leave-one-out cross-validation in terms of precision and recall via AdaL. The sizes of the points correspond to the numbers of predictions produced by a given combination. The color of the points corresponds to a clustering method.

**Supplementary Fig. S7.**
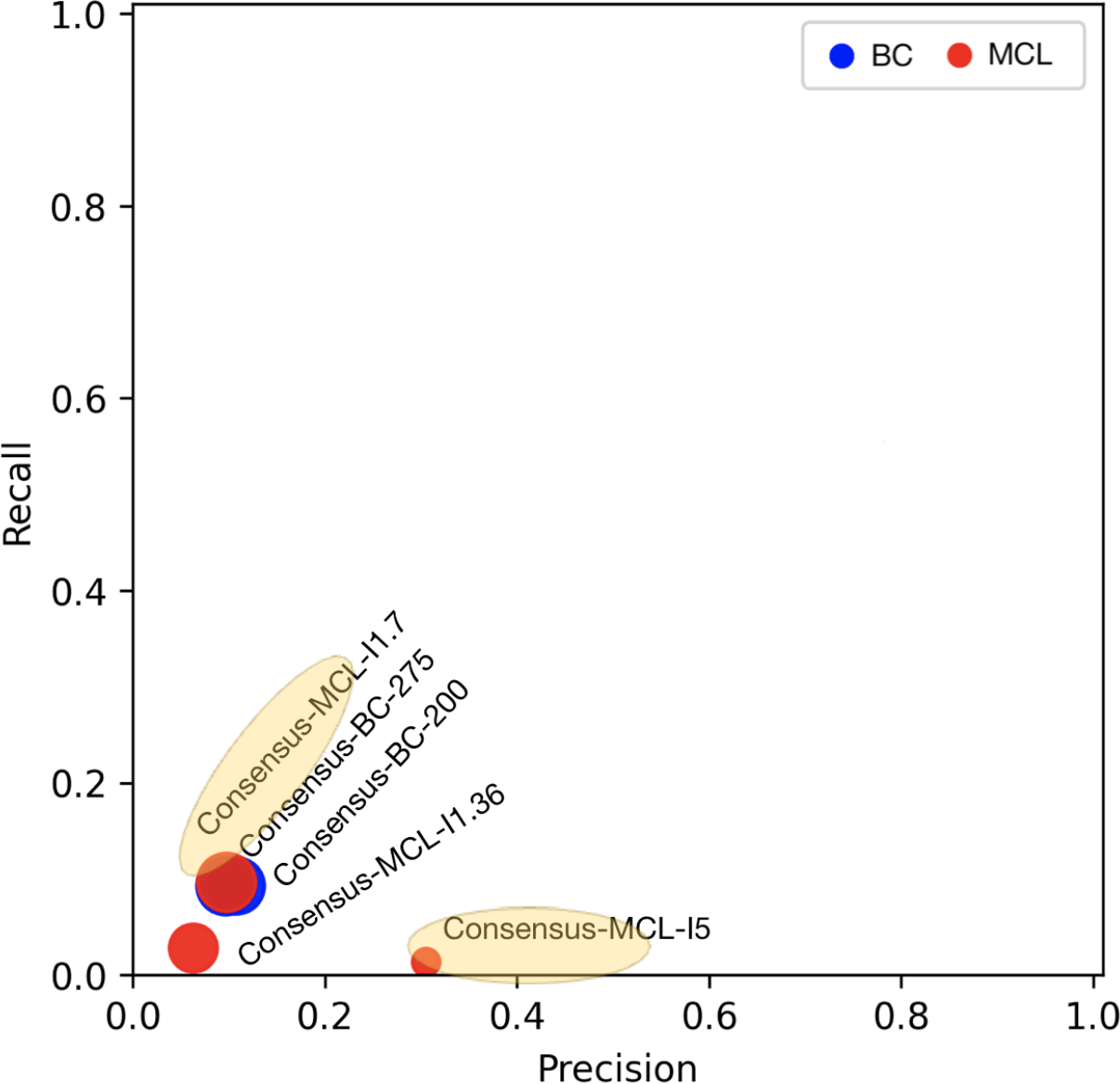
Accuracy of predicting gene-GO term associations in the leave-one-out cross-validation in terms of precision and recall via Consensus. The sizes of the points correspond to the numbers of predictions produced by a given combination. The color of the points corresponds to a clustering method.

**Supplementary Fig. S8.**
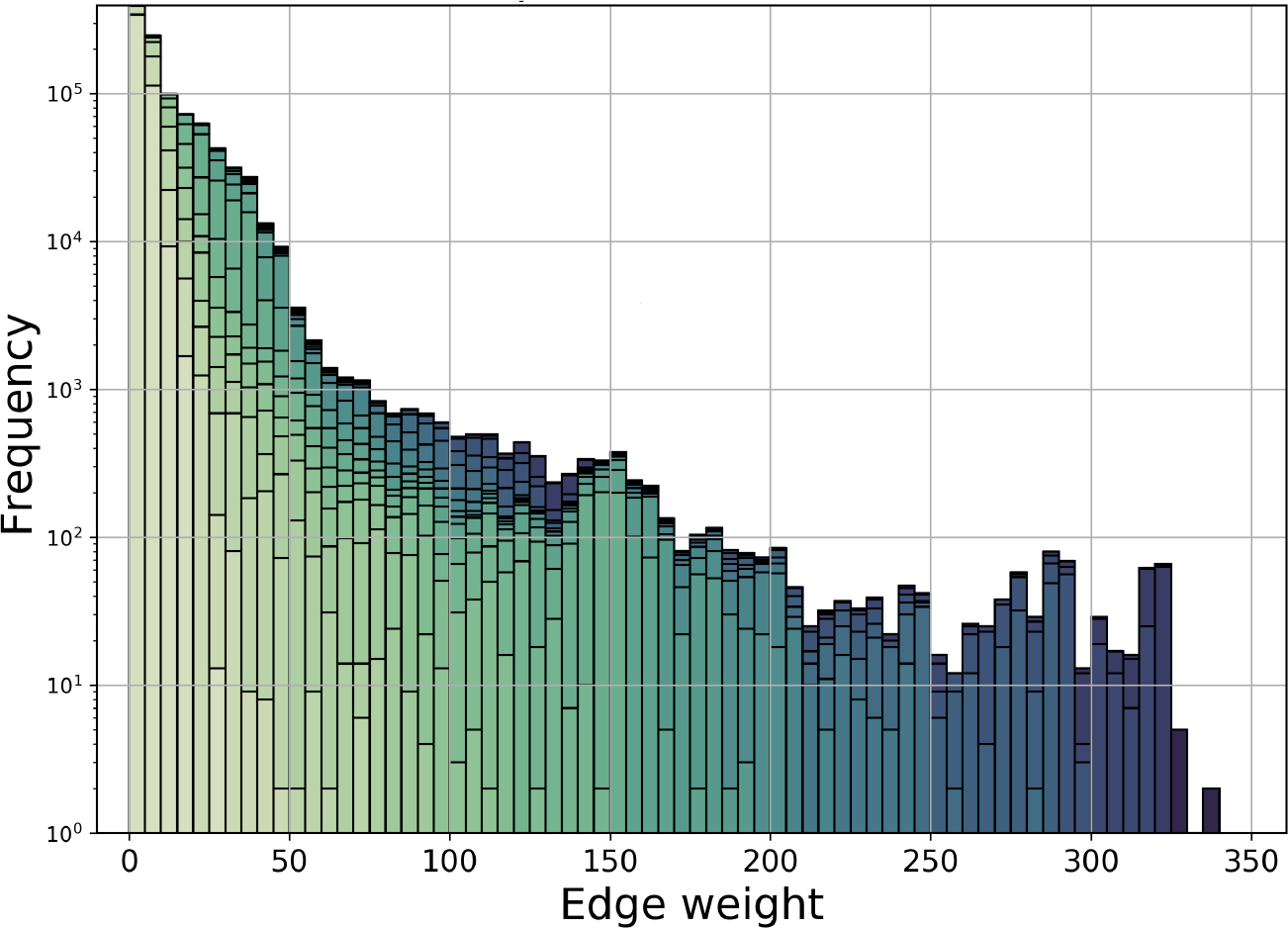
The distribution of confidence scores for predicted gene-gene interactions. The color shades represent the number of combinations of network and clustering method that support the corresponding association. The darker color the color, the higher the support. Analogous results for gene-GO term associations are shown in Figure 7 of the main paper.

## Conflict of interests

The authors declare that they have no conflict of interests.

## Author’s contributions

Q.L., K.B.S., M.F., and T.M. convinced and designed the study. Q.L. and K.B.S. collected and processed the (gene expression and gene-GO term annotation) data, with help from G.F. Q.L. performed most of the experiments (network inference, network clustering, gene-GO term annotation prediction, endocytosis analysis and interpretation), with help from K.B.S., M.S, and E.C. Q.L., K.B.S., M.F., and T.M. analyzed and discussed the results. Q.L. drafted the initial version of the paper, with significant contribution by K.B.S., M.F., and T.M. towards the final paper version. All authors approved the final version of the paper. T.M. supervised all computational aspects of the study and M.F. supervised all biological aspects of the study.

## Funding

This work is supported by NSF CAREER CCF-1452795 and National Institutes of Health grant P01 12072248 (MTF).

## Acknowledgements

We thank Khalique Newaz and Shawn Gu for their comments and suggestions for the clarity of our writings.

## Footnotes

1 https://www.genedb.org/

2 https://plasmodb.org/plasmo/app

3 https://www.genedb.org/

4 https://plasmodb.org/plasmo/app

5 https://micans.org/mcl/

## Notes

https://nd.edu/~cone/pfalGCEN/

### Competing Interest Statement

The authors have declared no competing interest.

### Summary of Updates

We have revisited and reorganized the writing for better readability.

## References

1. S. H. Adjalley, D. Scanfeld, E. Kozlowski, M. Llinas, and D. A. Fidock. Genome-wide transcriptome profiling reveals functional networks involving the plasmodium falciparum drug resistance transporters pfcrt and pfmdr1. BMC Genomics, 16:1090, 2015.

2. M. E. Ahsen, Y. Chun, A. Grishin, G. Grishina, G. Stolovitzky, G. Pandey, and S. Bunyavanich. NeTFactor, a framework for identifying transcriptional regulators of gene expression-based biomarkers. Scientific Reports, 9(1):1–13, 2019.

3. M. Bansal, V. Belcastro, A. Ambesi-Impiombato, and D. Di Bernardo. How to infer gene networks from expression profiles. Molecular Systems Biology, 3(1):78, 2007.

4. A.-L. Barabási. Network science. Philosophical Transactions of the Royal Society A: Mathematical, Physical and Engineering Sciences, 371(1987):20120375, 2013.

5. A. Bauer-Mehren, M. Bundschus, M. Rautschka, M. A. Mayer, F. Sanz, and L. I. Furlong. Gene-disease network analysis reveals functional modules in mendelian, complex and environmental diseases. PLOS ONE, 6(6):e20284, 2011.

6. J. Benesty, J. Chen, Y. Huang, and I. Cohen. Pearson correlation coefficient. In Noise Reduction in Speech Processing, pages 1–4. Springer, 2009.

7. J. Birnbaum, S. Scharf, S. Schmidt, E. Jonscher, W. A. M. Hoeijmakers, S. Flemming, C. G. Toenhake, M. Schmitt, R. Sabitzki, B. Bergmann, et al. A Kelch13-defined endocytosis pathway mediates artemisinin resistance in malaria parasites. Science, 367(6473):51–59, 2020.

8. S. v. Buuren and K. Groothuis-Oudshoorn. MICE: Multivariate imputation by chained equations in R. Journal of Statistical Software, pages 1–68, 2010.

9. D. Camacho, A. De La Fuente, and P. Mendes. The origin of correlations in metabolomics data. Metabolomics, 1(1):53–63, 2005.

10. E. J. Candés and B. Recht. Exact matrix completion via convex optimization. Foundations of Computational Mathematics, 9(6):717, 2009.

11. S. B. Carey, J. Jenkins, J. T. Lovell, F. Maumus, A. Sreedasyam, A. C. Payton, S. Shu, G. P. Tiley, N. Fernandez-Pozo, A. Healey, et al. Gene-rich UV sex chromosomes harbor conserved regulators of sexual development. Science Advances, 7(27):eabh2488, 2021.

12. A. Dobra, C. Hans, B. Jones, J. R. Nevins, G. Yao, and M. West. Sparse graphical models for exploring gene expression data. Journal of Multivariate Analysis, 90(1):196–212, 2004.

13. A. M. Dondorp, F. Nosten, P. Yi, D. Das, A. P. Phyo, J. Tarning, K. M. Lwin, F. Ariey, W. Hanpithakpong, S. J. Lee, et al. Artemisinin resistance in Plasmodium falciparum malaria. New England Journal of Medicine, 361(5):455–467, 2009.

14. A. J. Enright, S. Van Dongen, and C. A. Ouzounis. An efficient algorithm for large-scale detection of protein families. Nucleic Acids Research, 30(7):1575–1584, 2002.

15. A. Gorovits, E. Gujral, E. E. Papalexakis, and P. Bogdanov. Larc: Learning activity-regularized overlapping communities across time. In Proceedings of the 24th ACM SIGKDD International Conference on Knowledge Discovery & Data Mining, pages 1465–1474, 2018.

16. B. Greenwood and T. Mutabingwa. Malaria in 2002, 2002.

17. F. Hahne, W. Huber, R. Gentleman, S. Falcon, S. Falcon, and R. Gentleman. Hypergeometric testing used for gene set enrichment analysis. Bioconductor case studies, pages 207–220, 2008.

18. A. U. Hain, A. S. Miller, J. Levitskaya, and J. Bosch. Virtual screening and experimental validation identify novel inhibitors of the Plasmodium falciparum Atg8-Atg3 protein-protein interaction. ChemMedChem, 11(8):900–10, 2016.

19. T. Hastie, R. Mazumder, J. D. Lee, and R. Zadeh. Matrix completion and low-rank SVD via fast alternating least squares. The Journal of Machine Learning Research, 16(1):3367–3402, 2015.

20. R. C. Henrici, R. L. Edwards, M. Zoltner, D. A. van Schalkwyk, M. N. Hart, F. Mohring, R. W. Moon, S. D. Nofal, A. Patel, C. Flueck, et al. The Plasmodium falciparum artemisinin susceptibility-associated AP-2 adaptin μ subunit is clathrin independent and essential for schizont maturation. Mbio, 11(1):e02918–19, 2020.

21. G. Hu, A. Cabrera, M. Kono, S. Mok, B. K. Chaal, S. Haase, K. Engelberg, S. Cheemadan, T. Spielmann, P. R. Preiser, et al. Transcriptional profiling of growth perturbations of the human malaria parasite Plasmodium falciparum. Nature Biotechnology, 28(1):91–98, 2010.

22. G. Hu, M. Llinás, J. Li, P. R. Preiser, and Z. Bozdech. Selection of long oligonucleotides for gene expression microarrays using weighted rank-sum strategy. BMC Bioinformatics, 8:350, 2007.

23. P. Hunt, A. Martinelli, K. Modrzynska, S. Borges, A. Creasey, L. Rodrigues, D. Beraldi, L. Loewe, R. Fawcett, S. Kumar, et al. Experimental evolution, genetic analysis and genome re-sequencing reveal the mutation conferring artemisinin resistance in an isogenic lineage of malaria parasites. BMC Genomics, 11(1):1–13, 2010.

24. V. A. Huynh-Thu, A. Irrthum, L. Wehenkel, and P. Geurts. Inferring regulatory networks from expression data using tree-based methods. PLOS ONE, 5(9):1–10, 2010.

25. M. M. Ippolito, K. A. Moser, J.-B. B. Kabuya, C. Cunningham, and J. J. Juliano. Antimalarial drug resistance and implications for the who global technical strategy. Current Epidemiology Reports, 8:46–62, 2021.

26. J. Iqbal, M. Al-Awadhi, and S. Ahmad. Decreasing trend of imported malaria cases but increasing influx of mixed P. falciparum and P. vivax infections in malaria-free Kuwait. PLOS ONE, 15(12):e0243617, 2020.

27. S. Jahagirdar, M. Suarez-Diez, and E. Saccenti. Simulation and reconstruction of metabolite–metabolite association networks using a metabolic dynamic model and correlation based algorithms. Journal of Proteome Research, 18(3):1099–1113, 2019.

28. K. J. Karczewski, L. C. Francioli, G. Tiao, B. B. Cummings, J. Alföldi, Q. Wang, R. L. Collins, K. M. Laricchia, A. Ganna, D. P. Birnbaum, et al. The mutational constraint spectrum quantified from variation in 141,456 humans. Nature, 581(7809):434–443, 2020.

29. H. Kim, G. H. Golub, and H. Park. Missing value estimation for DNA microarray gene expression data: local least squares imputation. Bioinformatics, 21(2):187–198, 2005.

30. D. Kochar, A. Das, A. Kochar, S. Middha, J. Acharya, G. Tanwar, D. Pakalapati Subudhi, P. Boopathi, S. Garg, et al. A prospective study on adult patients of severe malaria caused by Plasmodium falciparum, Plasmodium vivax and mixed infection from Bikaner, northwest India. Journal of Vector Borne Diseases, 51(3):200, 2014.

31. P. Koskinen, P. Törönen, J. Nokso-Koivisto, and L. Holm. PANNZER: high-throughput functional annotation of uncharacterized proteins in an error-prone environment. Bioinformatics, 31(10):1544–1552, 2015.

32. N. Krämer, J. Schäfer, and A.-L. Boulesteix. Regularized estimation of large-scale gene association networks using graphical Gaussian models. BMC Bioinformatics, 10(1):384, 2009.

33. A. Lachmann, F. M. Giorgi, G. Lopez, and A. Califano. ARACNe-AP: gene network reverse engineering through adaptive partitioning inference of mutual information. Bioinformatics, 32(14):2233–2235, 2016.

34. D. J. LaCount, M. Vignali, R. Chettier, A. Phansalkar, R. Bell, J. R. Hesselberth, L. W. Schoenfeld, I. Ota, S. Sahasrabudhe, C. Kurschner, et al. A protein interaction network of the malaria parasite Plasmodium falciparum. Nature, 438(7064):103–107, 2005.

35. K. A. Lawson, C. M. Sousa, X. Zhang, E. Kim, R. Akthar, J. J. Caumanns, Y. Yao, N. Mikolajewicz, C. Ross, K. R. Brown, et al. Functional genomic landscape of cancer-intrinsic evasion of killing by t cells. Nature, 586(7827):120–126, 2020.

36. J. Le Bras and R. Durand. The mechanisms of resistance to antimalarial drugs in Plasmodium falciparum. Fundamental & Clinical Pharmacology, 17(2):147–153, 2003.

37. H. Li and J. Gui. Gradient directed regularization for sparse gaussian concentration graphs, with applications to inference of genetic networks. Biostatistics, 7(2):302–317, 2006.

38. Q. Li and T. Milenkovic. Supervised prediction of aging-related genes from a context-specific protein interaction subnetwork. IEEE/ACM Transactions on Computational Biology and Bioinformatics, 2021.

39. Q. Li, K. Newaz, and T. Milenković. Improved supervised prediction of aging-related genes via weighted dynamic network analysis. BMC bioinformatics, 22(1):1–26, 2021.

40. Q. Liao, C. Liu, X. Yuan, S. Kang, R. Miao, H. Xiao, G. Zhao, H. Luo, D. Bu, H. Zhao, et al. Large-scale prediction of long non-coding RNA functions in a coding–non-coding gene co-expression network. Nucleic Acids Research, 39(9):3864–3878, 2011.

41. K. Lu, K. Yang, E. Niyongabo, Z. Shu, J. Wang, K. Chang, Q. Zou, J. Jiang, C. Jia, B. Liu, et al. Integrated network analysis of symptom clusters across disease conditions. Journal of Biomedical Informatics, 107:103482, 2020.

42. L. Manning, M. Laman, I. Law, C. Bona, S. Aipit, D. Teine, J. Warrell Rosanas-Urgell, E. Lin, B. Kiniboro, et al. Features and prognosis of severe malaria caused by Plasmodium falciparum, Plasmodium vivax and mixed Plasmodium species in Papua New Guinean children. PLOS ONE, 6(12):e29203, 2011.

43. D. Marbach, J. C. Costello, R. Kuffner, N. M. Vega, R. J. Prill, D. M. Camacho, K. R. Allison, R. Bonneau, Y. Chen, J. J. Collins, F. Coordero, J. C. Costello, M. Crane, F. Dondelinger, M. Drton, R. Espositoo, R. Foygel, T. D. Consortium, and G. Stolvitzky. Wisdom of crowds for robust gene network inference. Nature Methods, 9(8):796–804, 2012.

44. A. A. Margolin, I. Nemenman, K. Basso, C. Wiggins, G. Stolovitzky, R. Dalla Favera, and Califano. ARACNE: an algorithm for the reconstruction of gene regulatory networks in a mammalian cellular context. In BMC Bioinformatics, volume 7, pages S1–S7. Springer, 2006.

45. R. Mazumder, T. Hastie, and R. Tibshirani. Spectral regularization algorithms for learning large incomplete matrices. The Journal of Machine Learning Research, 11:2287–2322, 2010.

46. R. A. C. Montes, G. Coello, K. L. González-Aguilera, N. Marsch-Martínez, S. de Folter, and E. R. Alvarez-Buylla. ARACNe-based inference, using curated microarray data, of arabidopsis thaliana root transcriptional regulatory networks. BMC Plant Biology, 14(1):97, 2014.

47. S. Nelson-Sathi, F. L. Sousa, M. Roettger, N. Lozada-Chávez, T. Thiergart, A. Janssen, D. Bryant, G. Landan, P. Schönheit, B. Siebers, et al. Origins of major archaeal clades correspond to gene acquisitions from bacteria. Nature, 517(7532):77–80, 2015.

48. P. K.-S. Ng, J. Li, K. J. Jeong, S. Shao, H. Chen, Y. H. Tsang, S. Sengupta, Z. Wang, V. H. Bhavana, R. Tran, et al. Systematic functional annotation of somatic mutations in cancer. Cancer Cell, 33(3):450–462, 2018.

49. J. Oberstaller, T. D. Otto, J. C. Rayner, and J. H. Adams. Essential genes of the parasitic apicomplexa. Trends in Parasitology, 37(4):304–316, 2021.

50. J. Oliveira-Ferreira, M. V. Lacerda, P. Brasil, J. L. Ladislau, P. L. Tauil, and C. T. Daniel-Ribeiro. Malaria in Brazil: an overview. Malaria Journal, 9(1):1–15, 2010.

51. W. H. Organization. Malaria. https://www.who.int/news-room/fact-sheets/detail/malaria, 2020.

52. W. H. Organization et al. The potential impact of health service disruptions on the burden of malaria: a modelling analysis for countries in sub-Saharan Africa. WHO, 2020.

53. M. Pham, S. Wilson, H. Govindarajan, C.-H. Lin, and O. Lichtarge. Discovery of disease-and drug-specific pathways through community structures of a literature network. Bioinformatics, 36(6):1881–1888, 2020.

54. M. S. Pieperhoff, M. Schmitt, D. J. Ferguson, and M. Meissner. The role of clathrin in post-Golgi trafficking in Toxoplasma gondii. PLOS ONE, 8(10):e77620, 2013.

55. A. Ramaprasad, A. Pain, and T. Ravasi. Defining the protein interaction network of human malaria parasite plasmodium falciparum. Genomics, 99(2):69–75, 2012.

56. A. K. Rider, T. Milenković, G. H. Siwo, R. S. Pinapati, S. J. Emrich, M. T. Ferdig, and N. V. Chawla. Networks’ characteristics are important for systems biology. Network Science, 2(2):139–161, 2014.

57. T. Rito, Z. Wang, C. M. Deane, and G. Reinert. How threshold behaviour affects the use of subgraphs for network comparison. Bioinformatics, 26(18):i611–i617, 2010.

58. S. S. Rund, B. Yoo, C. Alam, T. Green, M. T. Stephens, E. Zeng, G. F. George, A. D. Sheppard, G. E. Duffield, T. Milenković, et al. Genome-wide profiling of 24 hr diel rhythmicity in the water flea, Daphnia pulex: network analysis reveals rhythmic gene expression and enhances functional gene annotation. BMC Genomics, 17(1):653, 2016.

59. J. Schäfer and K. Strimmer. A shrinkage approach to large-scale covariance matrix estimation and implications for functional genomics. Statistical Applications in Genetics and Molecular Biology, 4(1), 2005.

60. G. H. Siwo, A. Tan, K. A. Button-Simons, U. Samarakoon, L. A. Checkley, R. S. Pinapati, and M. T. Ferdig. Predicting functional and regulatory divergence of a drug resistance transporter gene in the human malaria parasite. BMC Genomics, 16(1):1–13, 2015.

61. T. Spielmann, S. Gras, R. Sabitzki, and M. Meissner. Endocytosis in Plasmodium and Toxoplasma parasites. Trends in Parasitology, 36(6):520–532, 2020.

62. W. Stacklies, H. Redestig, M. Scholz, D. Walther, and J. Selbig. PCAMethods—a bioconductor package providing PCA methods for incomplete data. Bioinformatics, 23(9):1164–1167, 2007.

63. F. Supek, B. Matko, N. Skunca, and T. Smuc. REVIGO summarizes and visualizes long lists of Gene Ontology terms. PLOS ONE, 6(7):e21800, 2011.

64. D. Szklarczyk, A. L. Gable, K. C. Nastou, D. Lyon, R. Kirsch, S. Pyysalo, N. T. Doncheva, M. Legeay, T. Fang, P. Bork, et al. The STRING database in 2021: customizable protein–protein networks, and functional characterization of user-uploaded gene/measurement sets. Nucleic Acids Research, 49(D1):D605–D612, 2021.

65. J. Talapko, I. Škrlec, T. Alebić, M. Jukić, and A. Včev. Malaria: the past and the present. Microorganisms, 7(6):179, 2019.

66. Q. W. Tan and M. Mutwil. Malaria.tools—comparative genomic and transcriptomic database for Plasmodium species. Nucleic Acids Research, 48(D1):D768–D775, 08 2019.

67. N. Tangpukdee, C. Duangdee, P. Wilairatana, and S. Krudsood. Malaria diagnosis: a brief review. The Korean Journal of Parasitology, 47(2):93, 2009.

68. V. Thakur, M. Asad, S. Jain, M. E. Hossain, A. Gupta, I. Kaur, S. Rathore, S. Ali, N. J. Khan, and A. Mohmmed. Eps15 homology domain containing protein of Plasmodium falciparum (pfehd) associates with endocytosis and vesicular trafficking towards neutral lipid storage site. Biochimica et Biophysica Acta (BBA)-Molecular Cell Research, 1853(11):2856–2869, 2015.

69. O. Troyanskaya, M. Cantor, G. Sherlock, P. Brown, T. Hastie, R. Tibshirani, D. Botstein, and R. B. Altman. Missing value estimation methods for DNA microarrays. Bioinformatics, 17(6):520–525, 2001.

70. S. van Dam, U. Vosa, A. van der Graaf, L. Franke, and J. P. de Magalhaes. Gene co-expression analysis for functional classification and gene–disease predictions. Briefings in Bioinformatics, 19(4):575–592, 2018.

71. Y. Wang, A. Batra, H. Schulenburg, and T. Dagan. Gene sharing among plasmids and chromosomes reveals barriers for antibiotic resistance gene transfer. Philosophical Transactions of the Royal Society B, 377(1842):20200467, 2021.

72. Y. Wang, H. Tang, J. D. DeBarry, X. Tan, J. Li, X. Wang, T.-h. Lee, H. Jin, B. Marler, H. Guo, et al. MCScanX: a toolkit for detection and evolutionary analysis of gene synteny and collinearity. Nucleic Acids Research, 40(7):e49–e49, 2012.

73. M. T. Weirauch. Gene coexpression networks for the analysis of DNA microarray data. Applied Statistics for Network Biology: Methods in Systems Biology, 1:215–250, 2011.

74. D. J. Weiss, T. C. Lucas, M. Nguyen, A. K. Nandi, D. Bisanzio, K. E. Battle, E. Cameron, K. A. Twohig, D. A. Pfeffer, J. A. Rozier, et al. Mapping the global prevalence, incidence, and mortality of plasmodium falciparum, 2000–17: a spatial and temporal modelling study. The Lancet, 394(10195):322–331, 2019.

75. J. Whittaker. Graphical Models in Applied Multivariate Statistics. Wiley Publishing, 2009.

76. D. C. Wong, C. Sweetman, and C. M. Ford. Annotation of gene function in citrus using gene expression information and co-expression networks. BMC Plant Biology, 14(1):186, 2014.

77. D. Yang, Y. He, B. Wu, Y. Deng, M. Li, Q. Yang, L. Huang, Y. Cao, and Y. Liu. Drinking water and sanitation conditions are associated with the risk of malaria among children under five years old in sub-Saharan Africa: a logistic regression model analysis of national survey data. Journal of Advanced Research, 21:1–13, 2020.

78. J. Yang and J. Leskovec. Overlapping community detection at scale: a nonnegative matrix factorization approach. In Proceedings of the sixth ACM International Conference on Web Search and Data Mining, pages 587–596, 2013.

79. F.-D. Yu, S.-Y. Yang, Y.-Y. Li, and W. Hu. Co-expression network with protein–protein interaction and transcription regulation in malaria parasite plasmodium falciparum. Gene, 518(1):7–16, 2013. Proceedings of the 23rd International Conference on Genome Informatics (GIW 2012).

